# High-density mapping of QTL controlling agronomically important traits in quinoa (*Chenopodium quinoa* Willd.)

**DOI:** 10.1101/2022.03.29.486179

**Authors:** Nathaly Maldonado-Taipe, Federico Barbier, Karl Schmid, Christian Jung, Nazgol Emrani

**Author notes:** Corresponding author Correspondence to: Dr. Nazgol Emrani Plant Breeding Institute Christian-Albrechts-University of Kiel Olshausenstrasse 40 D-24118 Kiel Germany.

## Abstract

Quinoa is a pseudocereal originating from the Andean regions. In spite of quinoa’s long cultivation history, genetic analysis of this crop is still in its infancy. We aimed to localize QTL contributing to the phenotypic variation of agronomically important traits. We crossed the Chilean accession PI-614889 and the Peruvian accession CHEN-109, which depicted significant differences in days to flowering, days to maturity, plant height, panicle length, thousand kernel weight (TKW), saponin content, and mildew susceptibility. We observed sizeable phenotypic variation across F_2_ plants and F_3_ families grown in the greenhouse and in the field, respectively. We used Skim-seq to genotype the F_2_ population and constructed a high-density genetic map with 133,923 SNPs. Fifteen QTL were found for ten traits. Two significant QTL, common in F_2_ and F_3_ generations, depicted pleiotropy for days to flowering, plant height, and TKW. The pleiotropic QTL harbored several putative candidate genes involved in photoperiod response and flowering time regulation. This study presents the first high-density genetic map of quinoa that incorporates QTL for several important agronomical traits. The pleiotropic loci can facilitate marker assisted selection in quinoa breeding programs.

**Key message:** Skim-sequencing enabled the construction a high-density genetic map (133,923 SNPs) and fifteen QTL were detected for ten agronomically important traits.

## Introduction

Quinoa (*Chenopodium quinoa* Willd.) is a pseudocereal native to the Andean region of South America. It is an allotetraploid species (2n = 4x = 36), with a genome size of 1.45-1.50 Gb (Jarvis et al. 2017). Quinoa is characterized by its broad genetic variation and adaptation to biotic and abiotic stresses. It exhibits resistance to insects and diseases and tolerance to frost, drought, and salinity. Furthermore, quinoa seeds have outstanding physicochemical, nutritional, and functional properties for human consumption. Due to these unique qualities, quinoa is considered an option to improve world food security (Alandia et al. 2021).

Quinoa cultivation has transcended continental boundaries and it is present in Europe, Africa, and Asia (Alandia et al. 2020). However, substantial breeding efforts are still needed to explore all quinoa qualities and to expand its cultivation worldwide. Quinoa breeding aims for short, non-branching plants with a compact panicle, as well as increased tolerance to abiotic and biotic stresses. Nevertheless, the main breeding objective in quinoa remains to be the development of high-yielding varieties and, in temperate regions and high latitudes of Europe, North America, and China, the adaptation to long-day conditions (Murphy et al. 2018; Patiranage et al. 2021). Thus, for breeding quinoa, a better understanding of the molecular regulation of flowering time and day-length responsiveness is essential since yield potential and local adaptation are largely determined by these processes.

In spite of being a domesticated crop, quinoa has not yet reached its full potential but molecular and genetic technologies may help change this situation (Alandia et al. 2021). In this scenario, QTL mapping is useful to understand the genetic basis of quantitative traits. The use of sequencing technologies and computational analysis has made QTL detection easier. In skim sequencing (Skim-seq), genomes are sequenced at low coverage and sequence variants are called after mapping to a reference genome. Later, imputation is performed based on genetic linkage. Due to the large size of linkage blocks, Skim-seq is a suitable method for genotyping F_2_ and F_3_ segregating populations (Golicz et al. 2015; Kumar et al. 2021).

To date, only a few *C. quinoa* linkage maps are available. The first quinoa linkage map was constructed using 216 SSR (simple sequence repeats) markers using a recombinant inbred line (RIL) population. The map consisted of 38 linkage groups (LGs) covering 913 cM (Jarvis et al. 2008). Another linkage map contained 14,178 SNPs (KASPar genotyping) mapped in two RIL populations. This map consisted of 29 LGs spanning 1,404 cM (Maughan et al. 2012). A recent linkage map by Jarvis et al. (2017) combined the map from Maughan et al. (2012) with two new linkage maps. The resulting map contains 6,403 markers on 18 LGs spanning 2,034 centimorgans (cM). A few studies have attempted to identify loci for agronomically important traits in quinoa so far. Cervantes and van Loo (2017) identified QTL for color, flowering time, and yield-related traits using a F_2_ population of 94 individuals from a cross between ‘Carina Red’ (bitter, dark seed) and ‘Atlas’ (non-bitter EU variety). They used a linkage map constructed with 1,076 SNPs and localized two major QTL, one for days to floral bud appearance on chromosome Cq6B, and another one for seed characters on chromosome Cq2B. In addition, a recent genome-wide association study with 2.9 million markers uncovered significant marker-trait associations for days to flowering, days to maturity, plant height and panicle length on chromosome Cq2A (Patirange et al. 2020).

## Methods

### Plant material and growth conditions

The Chilean quinoa accession PI-614889 (seed parent; seed code 171115) was crossed with the Peruvian inbred line CHEN-109 (pollen parent, seed code 170876) applying hot water emasculation (Emrani et al. 2020). The F_1_ plant was selfed to give rise to the F_2_ population (seed code: 190031). The F_3_ population (seed codes: 191203-191562) consisting of 334 families, was produced by selfing F_2_ plants (Table S1). A total number of 336 F_2_ individuals and 10 plants of each parent were grown in 13 cm pots from March to October 2019 in a greenhouse under long-day conditions in Kiel, Germany. Seeds were harvested from August to October 2019. Three hundred thirty-four F_3_ families and their parental lines were mechanically sown in a plant-to-row scheme in the field in 2020 (10.0°E 54.3°N, Achterwehr, Germany). One hundred fifty seeds were sown in two-meter single rows (one cm sowing depth) with 50 cm spacing between rows under a complete randomized block design with two blocks. Mechanical weeding was carried out four weeks after sowing using a row crop cultivator and hand weeding was performed five and seven weeks after sowing. Thinning was performed six weeks after sowing, aiming at 20 plants per row distanced at 10 cm.

**Table 1.** Results from the quinoa populations grown in the greenhouse (F_2_) and in the field (F_3_) and their parental lines. DTF: days to flowering, DTM: days to maturity, PH: plant height, PL: panicle length, PD: panicle density, TKW: thousand kernel weight, SW: seed weight per plant, SN: seed number per plant, SC: Saponin content, MS: Mildew susceptibility. NA: data not available due to late-maturity genotype.

| Character | Population/parent | Mean | Minimum | Maximum | Transgression (%) |
| --- | --- | --- | --- | --- | --- |
| DTF | CHEN-109 | 2019: 61±2<sup>a</sup><br>2020: 72±3<sup>a</sup> | - | - | - |
|  | PI-614889 | 2019: 76±1<sup>b</sup><br>2020: 96±4<sup>b</sup> | - | - | - |
|  | F<sub>2</sub> | 69±5 | 55 | 87 | 29.17 |
|  | F<sub>3</sub> | 82±9 | 67 | 77 | 14.63 |
| DTM | CHEN-109 | 2019: 177±7<sup>b</sup><br>2020: NA | - | - | - |
|  | PI-614889 | 2019: 110±2<sup>a</sup><br>2020: 129±4 | - | - | - |
|  | F<sub>2</sub> | 137±14 | 110 | 194 | 11.91 |
|  | F<sub>3</sub> | 152±9 | 123 | 165 | NA |
| PH (cm) | CHEN-109 | 2019: 151.40±7.06<sup>b</sup><br>2020: 164.74±27.56<sup>b</sup> | - | - | - |
|  | PI-614889 | 2019: 83.90±6.00<sup>a</sup><br>2020: 137.11±13.98<sup>a</sup> | - | - | - |
|  | F<sub>2</sub> | 136.3±20.63 | 83.00 | 193.00 | 22.02 |
|  | F<sub>3</sub> | 163.14±27.85 | 50.00 | 265.00 | 66.49 |
| PL (cm) | CHEN-109 | 2019: 37.20±4.32<sup>b</sup><br>2020: 32.37±6.74<sup>a</sup> | - | - | - |
|  | PI-614889 | 2019: 16.30±2.49<sup>a</sup><br>2020: 29.47±9.41<sup>a</sup> | - | - | - |
|  | F<sub>2</sub> | 24.07±6.14 | 14.00 | 50.00 | 8.33 |
|  | F<sub>3</sub> | 32.85±10.4 | 10.00 | 80.00 | 76.31 |
| TKW (g) | CHEN-109 | 2019: 2.00±0.14<sup>b</sup><br>2020: 1.52±0.06<sup>b</sup> | - | - | - |
|  | PI-614889 | 2019: 2.44±0.31<sup>a</sup><br>2020: 2.64±0.03<sup>a</sup> | - | - | - |
|  | F<sub>2</sub> | 2.75±0.49 | 1.18 | 3.91 | 77.97 |
|  | F<sub>3</sub> | 2.62±0.29 | 1.58 | 3.45 | 48.17 |
| SW (g) | CHEN-109 | 2019: 1.07±0.26<sup>b</sup> | - | - | - |
|  | PI-614889 | 2019: 5.90±0.88<sup>a</sup> | - | - | - |
|  | F<sub>2</sub> | 4.10±1.48 | 0.06 | 7.22 | 13.29 |
| SN | CHEN-109 | 2019: 633±161<sup>b</sup> | - | - | - |
|  | PI-614889 | 2019: 2,432±328<sup>a</sup> | - | - | - |
|  | F<sub>2</sub> | 1469±471 | 51 | 3,251 | 6.25 |
| SC | CHEN-109 | 2019: 6.84±0.20<sup>b</sup><br>2020: 6.5 ± 2.12<sup>b</sup> | - | - | - |
|  | PI-614889 | 2019: 3.20±1.09<sup>a</sup><br>2020: 2.00±0.00<sup>a</sup> | - | - | - |
|  | F<sub>2</sub> | 5.68±4.22 | 2.00 | 24.00 | 47.02 |
|  | F<sub>3</sub> | 12.61±8.28 | 2.00 | 30.00 | 57.97 |
| MS | CHEN-109 | 2020: 2.95±0.23<sup>b</sup> | - | - | - |
|  | PI-614889 | 2020: 2±0.64<sup>a</sup> | - | - | - |
|  | F<sub>3</sub> | 2.34±0.69 | 1 | 3 | 59.42 |

Different letters <sup>a,b</sup> indicate significant differences between parental lines in each year (F<sub>2</sub>:2019; F<sub>3</sub>:2020) (t-test, $\alpha=0.05$ ).

### Phenotypic evaluation

The following traits were assessed in both populations: days to flowering (DTF), days to maturity (DTM), plant height (PH), panicle length (PL), panicle density (PD), saponin content (SC), and thousand kernel weight (TKW) (Table S2). Mildew susceptibility (MS) was recorded only in the F_3_ population in the field, while seed weight per plant (SW) and seed number per plant (SN) were recorded only in the F_2_ population. In the F_2_ trial, 336 individual plants and 10 plants of each parental line were phenotyped. In the field, ten plants per block and family and the parental lines, were phenotyped (a total of 6,720 plants). Plants with significant biotic stress damage in the field (e.g., insect damage) were excluded from phenotyping (Table S1). We carried out phenotyping at different BBCH stages, which were defined by Sosa Zuniga et al. (2017). Additionally, in order to verify genetic segregation in the F_2_ generation, we phenotyped red axil pigmentation in all 336 individuals.

**Table 2.** Statistical parameters calculated for eight phenotypic characters measured in the F_3_ population. V_P_: phenotypic variance, V_G_: genotypic variance, V_E_: environmental variance, PCV: phenotypic coefficient of variation, GCV: genotypic coefficient of variation, ECV: environmental coefficient of variation, h^2^: broad sense heritability, GA: genetic advance as percentage of mean. DTF: days to flowering, DTM: days to maturity, PH: plant height, PL: panicle length, PD: panicle density, TKW: thousand kernel weight, SC: saponin content, MS: Mildew susceptibility.

| Parameter | DTF | DTM | PH | PL | PD | MS | TKW | SC |
| --- | --- | --- | --- | --- | --- | --- | --- | --- |
| V<sub>P</sub> | 72.54 | 81.01 | 775.68 | 108.15 | 1.10 | 0.48 | 0.02 | 8.28 |
| V<sub>G</sub> | 60.20 | 64.62 | 298.20 | 41.11 | 0.90 | 0.40 | 0.02 | 7.22 |
| V<sub>E</sub> | 12.34 | 16.39 | 477.49 | 67.03 | 0.20 | 0.08 | 0.002 | 1.06 |
| PCV (%) | 10.34 | 5.94 | 17.07 | 31.66 | 42.06 | 41.67 | 5.70 | 22.81 |
| GCV (%) | 9.42 | 5.30 | 10.59 | 19.52 | 38.12 | 37.93 | 5.44 | 21.30 |
| ECV (%) | 4.26 | 2.67 | 13.39 | 24.93 | 17.78 | 17.26 | 1.70 | 8.16 |
| h<sup>2</sup> (%) | 82.99 | 79.77 | 38.44 | 38.02 | 82.14 | 82.85 | 91.06 | 87.19 |
| GA (%) | 17.68 | 9.76 | 13.52 | 24.79 | 71.16 | 71.12 | 10.70 | 40.97 |

### DNA isolation and PCR

In order to verify genetic segregation by molecular marker analysis, leaf genomic DNA was isolated from 48 F_2_ plants and 194 F_3_ plants by the standard CTAB method. We used the InDel marker JASS5 (Fw: AGCCATTGCACTATGCCCTCTC; Rv: TGGCCCAACACCTAAGTGACG) (Zhang et al. 2017). Polymerase chain reaction (PCR) and agarose gel electrophoresis were carried out following the details presented in Table S3.

**Table 3.**
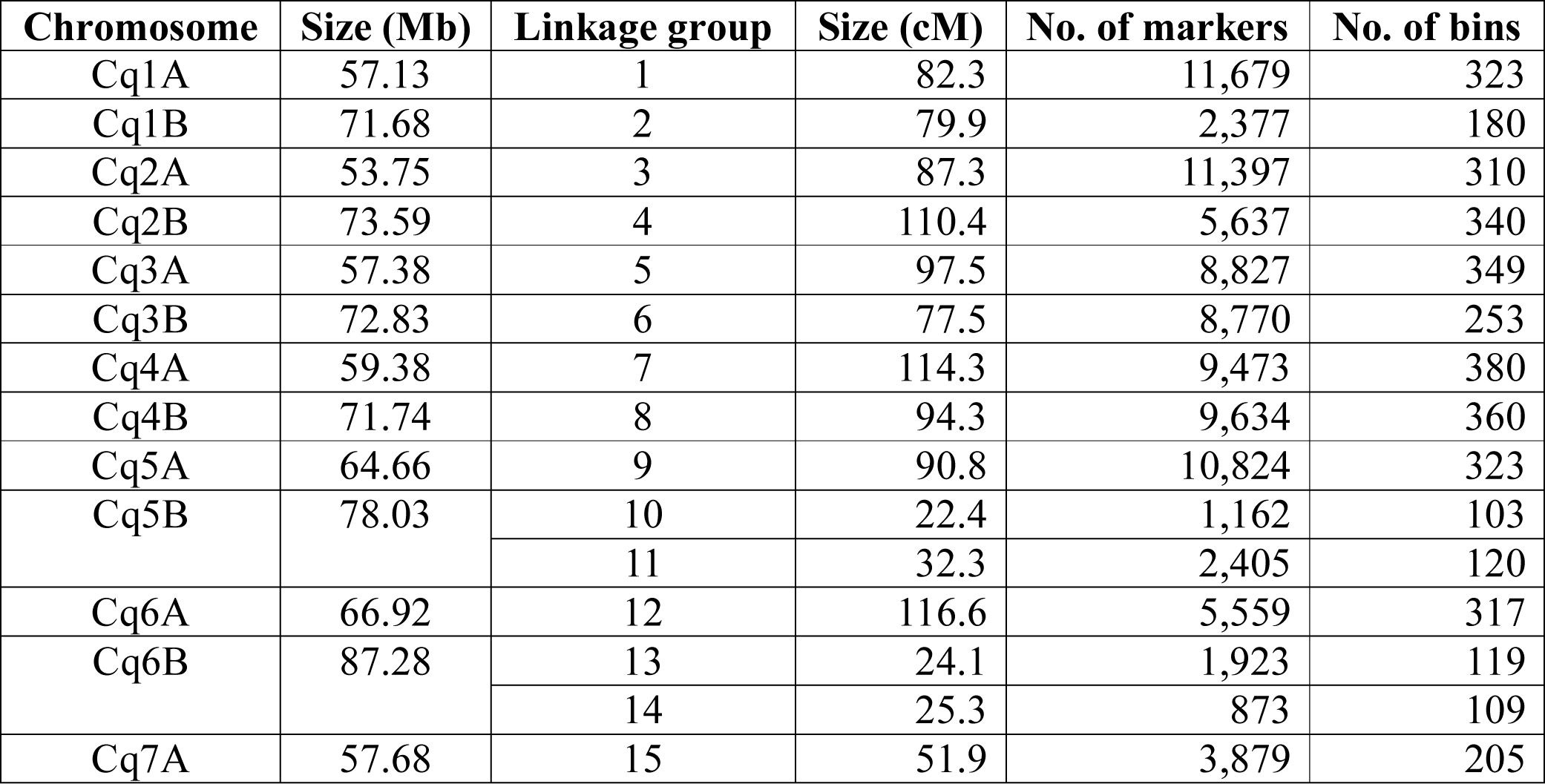
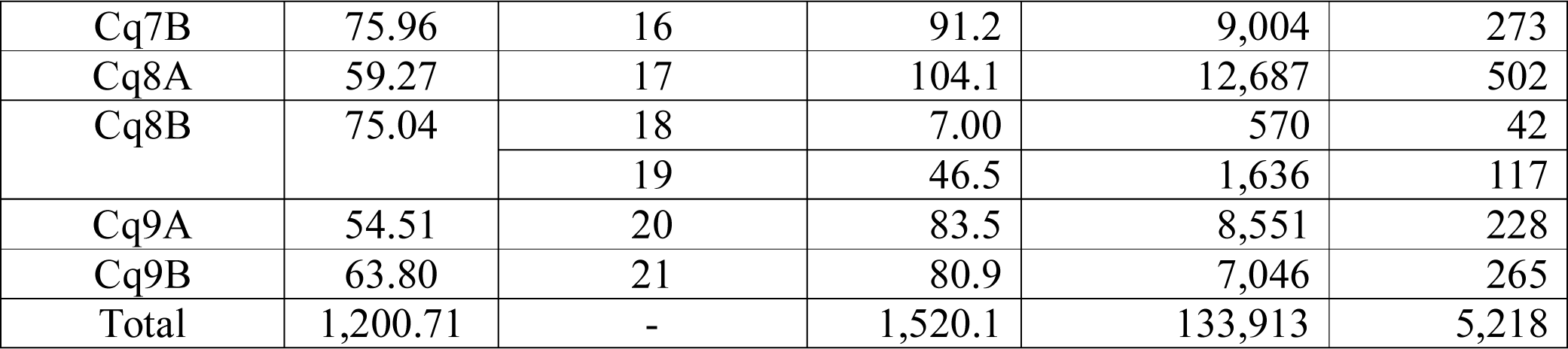
Summary statistics of the quinoa linkage map based on F_2_ plants derived from a cross between CHEN-109 and PI-614889. The physical size of each chromosome was taken from the reference genome QQ_74 (CoGe Genome ID: id60716).

For whole genome sequencing, we sampled young leaves from 336 F_2_ plants at BBCH 22 and freeze-dried them. We extracted DNA from these samples by a modified protocol of the Genomic Micro AX Blood Gravity kit (A&A Biotechnology, Gdansk, Poland). We verified the quality of the isolated DNA by agarose gel electrophoresis (0.8%, 60 V, 60 min).

### Whole genome sequencing and bioinformatics

Whole-genome sequencing libraries were constructed using the protocol of Baym et al. (2015) and normalized for equimolarity using a BioTec Synergy HTC multimode plate reader. The library was sequenced by Illumina NovaSeq PE150. We aimed to ∼1x coverage per sample of whole-genome sequences of the F_2_ individuals (Skim-seq). The genome sequences of both parents, CHEN-109 and PI-614889, were already available with a coverage of 7.45x and 8.00x, respectively (Patirange et al. 2020). We trimmed raw reads with Trim_galore v 0.6.4 (parameters -q 30 –fastqc –paired) (Krueger 2015), sorted and indexed them with SAMTOOLS 1.10 (Li et al. 2009) and deduplicated them with MarkDuplicates (parameter REMOVE_DUPLICATES = TRUE tool of PICARD v2.21.9) (Broad-Institute 2019). Quality control was done withFastQC and MultiQC by removing reads containing N > 10% (N represents the percentage of the nucleotides that cannot be determined) and a quality base filter of Qscore <= 5 (over 50% of the total base). We mapped the reads to the Quinoa Reference Genome QQ74_V2 (CoGe Genome ID: id60716). We called variants using HaplotypeCaller in -ERC GVCF mode (Poplin et al. 2017). Markers were named as “chromosome number_physical position” (e.g., chr12_ 2345937). We kept only homozygous loci within each parent and considered only SNPs with a minimum base quality of 30 (minQ=30) and minor allele frequency (maf): 0.1. Then, we imputed the missing data by FSFHap (maf: 0.1 MaxMissing: 0.8; Window: 50) (Swarts et al. 2014). After imputation, we applied the following filters: min-alleles: 2; max-alleles: 2 max-missing: 0.3; maf: 0.1. Finally, we transformed the data to a parent-based format (.abh) by using GenosToABH plugin from TASSEL (Glaubitz et al. 2014), using the codes A: pollen parent, B: seed parent, H: heterozygous. We performed quality control of the imputed data in .abh format by segregation distortion and percentage of missing data (ABHgenotypeR R package) (Reuscher and Furuta 2016). The bioinformatics pipeline is illustrated in Figure S1A.

### Linkage map construction

First, we performed final filtering of the F_2_ population (Figure S1B). We excluded F_2_ plants with more than 30% missing data. Only markers present in more than 302 F_2_ plants and fitting a 1:2:1 (α=0.05) segregation ratio were used for linkage studies. We also excluded “identical” individuals with >95% sequence similarity. Then, we constructed the genetic map by MSTMap (Wu et al. 2008) with the following parameters: Kosambi function, cut-off p-value = 1e-09, no_map_dist: 15, no_map_size: 2, missing_threshold: 0.1 (Supplementary File.map). Markers with an estimated genetic distance ≤ 1.00E-04 cM were clustered into bins. Finally, we performed several analyses for quality control of our genetic map. To begin with, we checked segregation distortion and estimated number of crossovers and double-crossover following the guidelines given by Broman (2010). In addition, we analyzed parental allele frequencies and collinearity of our linkage map with the physical map. Moreover, we used a heatmap of out linkage map to look for switched alleles. We used LinkageMapViewR, ASMapR, and R/qtl for quality control of the genetic map (Broman et al. 2003; Ouellette et al. 2018; Taylor and Butler 2017).

### QTL mapping and pleiotropy analysis

We carried out QTL mapping by composite interval mapping using the software package R/qtl version 1.46-2 (Haley-Knott with forward selection to three markers and a window size of 10 cM) (Broman and Sen 2009) (Supplementary Files .csv). The threshold for the logarithm of odds (LOD) for a significant QTL declaration was calculated by 1000 permutations of the genome-wide maximum LOD. The 95^th^ and 99^th^ percentile of this distribution were used as the genome wide LOD thresholds (5% and 1% LOD thresholds). The confidence intervals were calculated using the 95% Bayes credible interval method. QTL effects were calculated with the nearest markers as the phenotypic differences between marker genotypes. The percentage of phenotypic variation explained by each QTL (R²) was estimated by “drop-one-QTL-at-a-time” analysis. A simple additive model for multiple QTL was generated for each trait using the multiple imputation method and the Haley-Knott regression. When a putative pleiotropy was observed, it was confirmed by qtlpvl R package, and a multiple trait QTL analysis was performed (Tian et al. 2016). After confirmation of pleiotropy, pleiotropic sites were analyzed as single multitrait QTL (scanone.mvn) to obtain Bayes intervals and R² values.

### Epistasis analysis

A genome-wide epistasis analysis was performed to describe how alleles influence each other. For this purpose, we used cape R package (Tyler et al. 2013). To facilitate the analysis in terms of computational time, the software decomposed the phenotypes matrix into eigentraits (ET) by singular value decomposition (SVD). Then, we selected the two ET capturing the highest total variance among the traits to perform a pairwise scan of the variants (SNPs). From this scan, the software found interactions between alleles (epistasis), and the epistatic models were combined across ET to find allelic effects on the phenotypes included in the ETs. Positive and negative allelic effects refer to the reparametrized coefficient (either <0 or >0) from the pairwise regression as described by Tyler et al. (2013). Ultimately, the results of this analysis describe how alleles influence each other, in terms of enhancement (positive coefficient) and suppression (negative coefficient), as well as how gene variants influence phenotypes. The results of this analysis were plotted as heatmaps. ET contribution to the phenotypes was estimated for all bins of our genetic map, and heatmaps were constructed with 1,000 randomly selected markers. Effect calculations were performed in reference to the seed parent (PI-614889) allele.

### Candidate gene identification and haplotype analyses

We retrieved annotated genes from the reference genome within the regions of the confidence intervals of each QTL to explore possible candidate genes (.gff from QQ74_V2; CoGe Genome ID: id60716). We selected preliminary candidate genes using the UniProt Knowledgebase database (UniProtKB). A gene was considered a candidate when a related function to the identified QTL was already described in other plant species. Then, we searched for variants (SNPs and InDels) within the parental sequences. From this search, we kept homozygous genes and gave preference to those variants with a putative effect on the function of the encoded protein. Following, we evaluated haplotype of the selected variants in the F_2_ population, as follows: we clustered the phenotypes according to the corresponding genotype at the variant site; later, we performed t-tests (α=0.05) to compare DTF, PH, and TKW among the created clusters. To further evaluate the phenotypic effect of the variants, we used whole-genome sequencing and phenotypic data of 310 quinoa accessions grown in a two-year experiment in Kiel, Germany. This dataset comprises 2.9 million high confidence SNP and 414,891 InDel loci (Patirange et al. 2020). We followed the same procedure as for the haplotype evaluation in the F_2_ population. We assigned letters for each allele to describe the genotypes (e.g., *A_1_A_1_* homozygous, *A_1_A_2_* heterozygous). A complete description of the nomenclature is given in Table S4.

### Heritability estimates and statistical analysis

The phenotypic (V_P_), genotypic (V_G_), environmental variances (V_E_), and the broad sense heritabilities (h^2^) were estimated using F_3_ data (Falconer 1996). The heritability values were classified as low (below 30%), medium (30-60%), and high (above 60%) as suggested by Johnson et al. (1955). Genotypic coefficients of variation (GCV), phenotypic coefficients of variation (PCV), environmental coefficients of variation (ECV), and genetic advance with a selection intensity of 5% (GA) were calculated as described by Singh and Chaudhary (1977). In addition, phenotypic correlation coefficients (Pearson’s r) of quantitative traits within and between the F_2_ and F_3_ populations were estimated using the phenotypic value of each F_2_ plant and the average value of each F_3_ family.

The mean of each F_3_ family or parental line was used to estimate the corresponding phenotypic value. The percentage of transgression was defined as the number of plants with a phenotypic value exceeding the mean value of the superior parent or below the mean value of the inferior parent over the total number of phenotyped plants for the corresponding trait. We performed t-tests (α=0.05) to detect significant differences between phenotypic values of the parental lines.

## Results

### Segregation and phenotypic analysis of F_2_ and F_3_ populations

We verified the expected 1:2:1 genetic segregation in the F_2_ population by two approaches: phenotyping of the red axil pigmentation (complete dominance of red color over green color) (Simmonds 1971) and molecular marker analysis. We phenotyped red axil pigmentation in all 336 F_2_ individuals, while genotyping was carried out for 48 individuals. Likewise, the expected segregation in the F_3_ generation (3:2:3) was verified by molecular markers. One hundred ninety-four plants were genotyped from 20 randomly selected F_3_ families (Table S5 and Figure S2).

Both populations, F_2_ and F_3_, exhibited a vast phenotypic variation under field and greenhouse conditions (Figure S3). Moreover, substantial transgressive segregation was found for all traits (Table 1). The highest transgression percentage was found for TKW in the F_2_ generation. On the other hand, heritabilities ranged between 38.02 and 91.06% with TKW exhibiting the highest heritability value (91.06%). Besides, DTF, DTM, PD, and MS showed high heritability (79.77 to 82.99%) while PH and PL exhibited moderate heritability (38.44% and 38.02%, respectively) (Table 2). Importantly, only 34.9% of the plants reached maturity before harvesting in the field (October 2020), resulting in fewer plants being phenotyped for DTM in the F_3_ population (Table S1).

Then, we calculated correlations between all evaluated traits within years. The highest correlation was found between DTF and PH (Pearson’s r in F_2_: 0.69; Pearson’s r in F_3_: 0.63) (Fig 1). Both traits, PH and DTF, were significantly correlated with DTM. Furthermore, DTF showed a high negative correlation with TKW (F_2_ and F_3_) and with SN and SW (only F_2_). In general, taller plants flowered later, reached maturity later, and depicted a reduction in the yield traits values, while shorter plants flowered earlier, reached maturity earlier, and showed higher values for the yield-related traits. Additionally, significant correlations for DTF, DTM, PH, PD, and PL between the F_2_ plants grown under greenhouse conditions and their F_3_ progenies were calculated, with the highest values for DTF (0.73) and PH (0.66). Surprisingly, SC showed a low correlation between years (Pearson’s r: 0.17).

**Fig 1.**
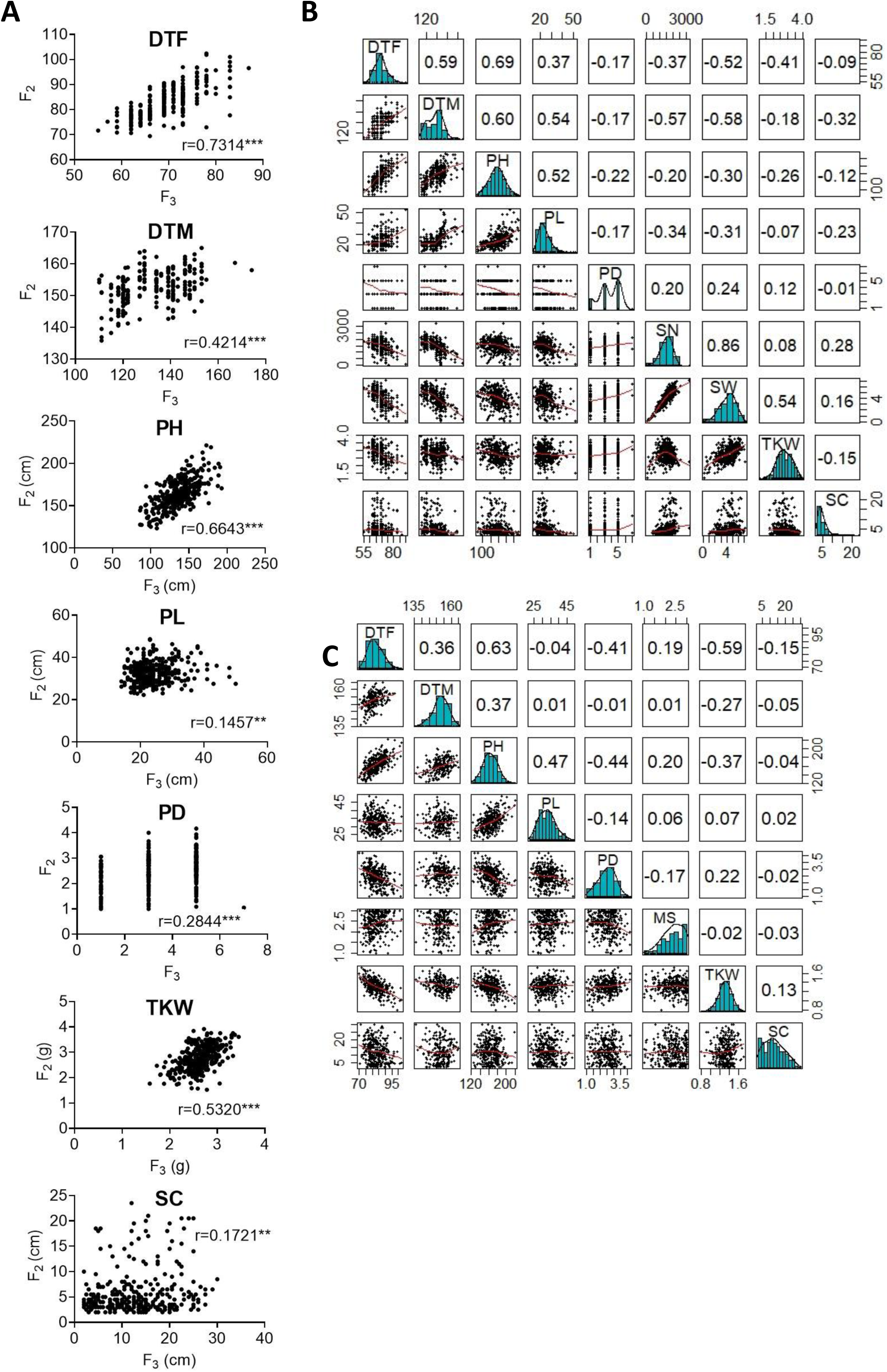
Correlation between phenotypic traits measured in the F_2_ and F_3_ populations. (A) Pearson’s r correlations between F_2_ and F_3_ populations. (B) Correlations between nine traits measured in the F_2_ population (C) Correlations between eight traits measured in the F_3_ population. In (B) and (C), bivariate scatter plots are shown below the diagonal, histograms on the diagonal, and the Pearson correlation above the diagonal. DTF: days to flowering, DTM: days to maturity, PH: plant height, PL: panicle length, PD: panicle density, TKW: thousand kernel weight, SW: seed weight per plant, SN: seed number per plant, MS: Mildew susceptibility, SC: saponin content

### Sequencing the F_2_ population revealed millions of SNPs

We sequenced the genomes of 336 F_2_ plants, the parents of the F_3_ families grown in the field. Skim-Seq by Illumina NovaSeq PE150 resulted in a total data output of 4.98 million raw PE reads on average per individual (∼1.07x coverage). All reads passed the quality base filter requirement (Qscore<= 5), and 0.0053% of the raw data were removed due to a high number of nucleotides that could not be determined (N>10%)

Seventeen million SNPs were obtained after mapping and variant calling (Figure S4). First, these SNPs were filtered by maf: 0.1 and minQ30, producing a data set of 4 million high-quality SNPs with a high percentage of missing data (∼86%) (Figure S5). After imputation, the proportion of missing data was reduced from ∼86.0% to ∼11.0%. Following the next filtering steps, we obtained a set of 249,744 high-quality biallelic SNPs with 4.2% missing data (Figure S6), 21.2% homozygous markers for the pollen parent allele, 26.5% homozygous markers for the seed parent allele, and 52.3% heterozygous markers (Fig 2). We used this set of markers for genetic map construction.

**Fig 2.**
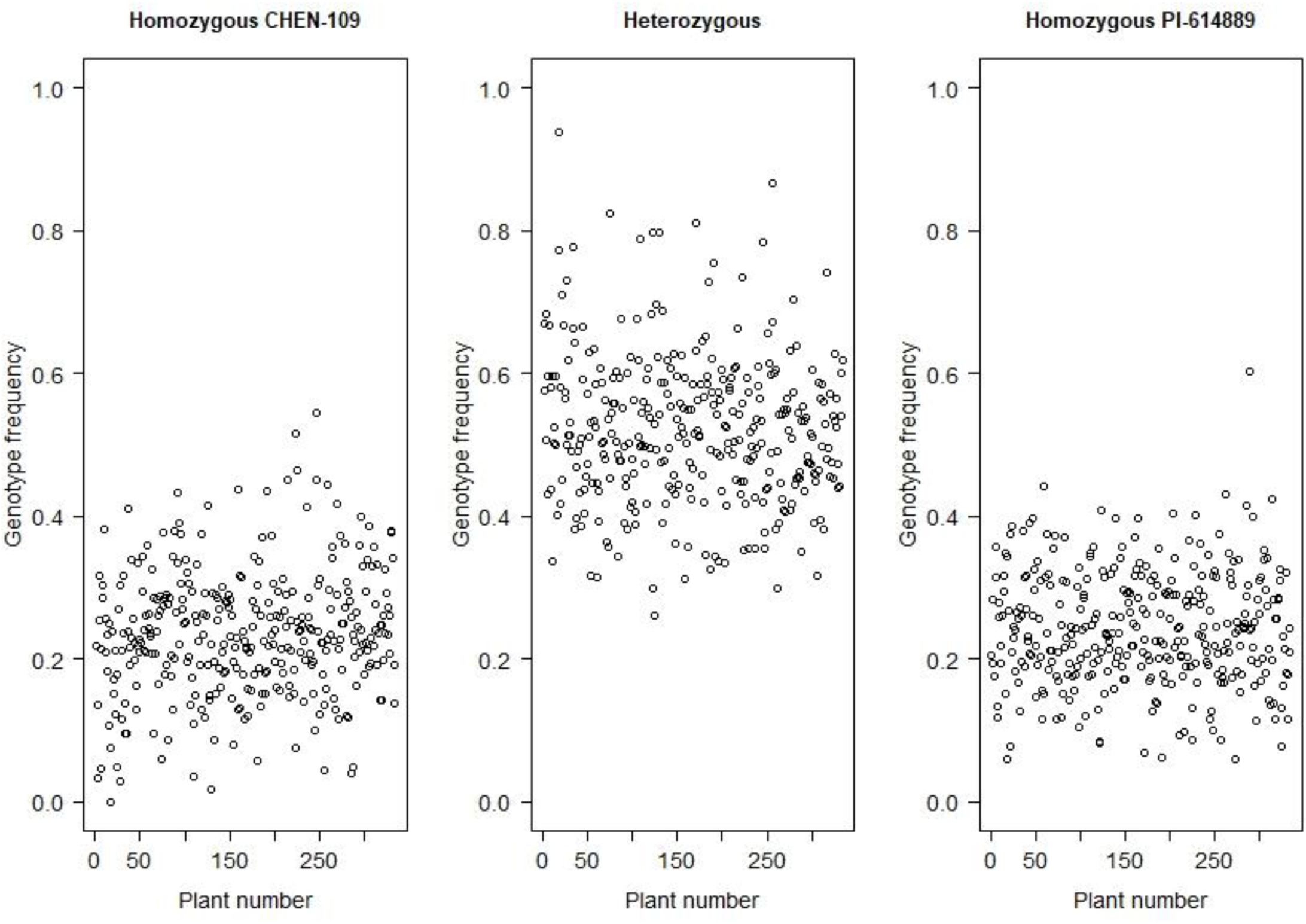
Frequency distribution of the homozygous genotype from the parent CHEN-109, the homozygous genotype from the parent PI-614889 and heterozygous genotype, for each of the F_2_ individuals

### Construction of a high-density linkage map

Ahead of genetic map construction, the F_2_ population sequences were cleaned anew based on our quality criteria. Two plants were removed due to > 30% missing data (Figure S6A), and 15,933 markers were removed because they were missing from >10% of the population (Figure S6B). Another 99,898 markers were removed because they did not segregate in the expected 1:2:1 manner. No plants had to be removed due to high (>95%) sequence similarity to another F_2_ plant (Figure S6C). As outcome of our final filtering, 334 F_2_ plants were used to construct a genetic map with 133,913 markers, resulting in an average density of one marker per ∼8.97 Kb. The resulting genetic map consists of 21 linkage groups (LG), with the chromosomes 5B, 6B and 8B split into two LGs each (Table 3, Figure S7). Moreover, the linkage map has an average density of ∼ 88 markers per cM, where one cM corresponds to ca. 0.83 Mb (Figure S8). For further steps, we created 5,219 bins where the markers with a genetic distance ≤ 1.00E-04 cM were clustered into.

To continue, we carried out several quality control analyses on the genetic map. First, we checked the number of single and double crossovers per plant, which ranged from 10 to 105 and 0 to 9, respectively (Figure S9A). We did not find any outlier plants depicting a significant higher number of crossovers and double crossovers than the ones observed for the population, which would have indicated potential genotyping errors (Figure S9B). Second, we analyzed the collinearity of our genetic map with the physical map from the reference genome sequence V2 and observed a high collinearity. We observed major gaps at the centromeres and an inversion at LG 7 (Fig 3 and Figure S7). Third, we investigated switched alleles by a heatmap (Fig 4). We did not find switched alleles, which would be indicated by pairs of markers with low LOD scores and low recombination fraction. Moreover, we inspected the parental allele frequencies in each linkage group, which were as expected: 0.25 of CHEN-109 genotype, 0.25 of PI-614889 genotype, and 0.5 heterozygous genotypes (Figure S10).

**Fig 3.**
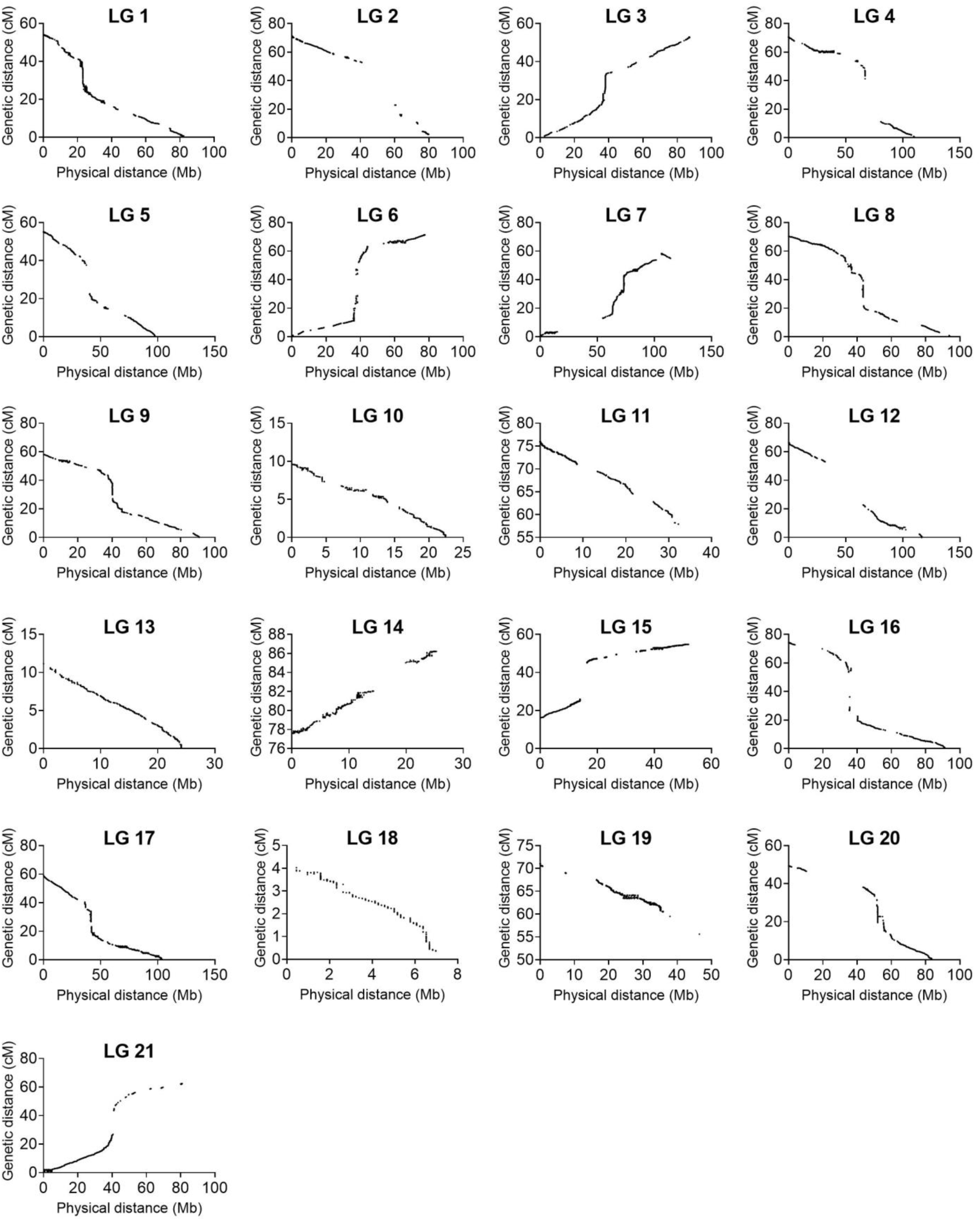
Collinearity between the linkage map constructed in this study and the physical map from the reference genome QQ74_V2 (CoGe Genome ID: id60716). The graphs were constructed with 133,913 non-binned markers.

**Fig 4.**
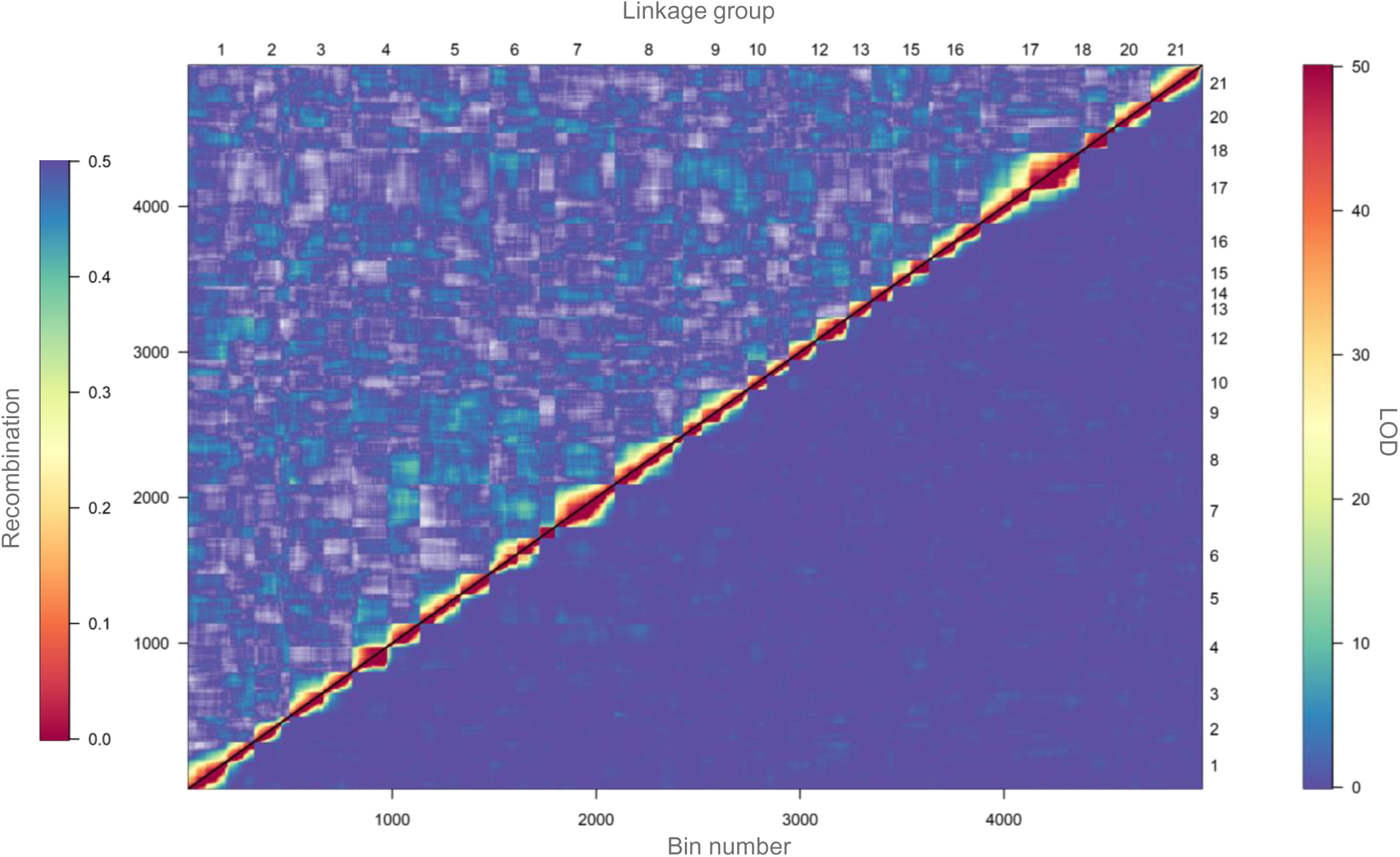
Heatmap of pairwise recombination fractions and LOD scores based on 5,219 bins. Estimated recombination fractions between binned-markers are shown above the diagonal and LOD scores below the diagonal. Red colors indicate closely linked binned-markers (high LOD score and low recombination fraction) whereas blue colors indicate non-linked binned-markers (low LOD score and high recombination fraction). A LOD score of 50 corresponds to a recombination fraction of zero.

### QTL mapping, pleiotropic loci identification, and epistasis calculation

We mapped QTL for ten agronomically important traits using phenotypic data of 334 F_2_ plants and 328 F_3_ families, which had passed our quality check (Figure S6). Fifteen QTL were identified, ranging from one to three QTL per trait (Table 4). We found pleiotropy at seven QTL, which were named with the prefix “*pleio*” (Fig 5 and Figure S11). Two QTL (*pleio4.1* and *pleio14.1*) were in common between F_2_ and F_3_, whereas six and eight QTL were found only in F_2_ or F_3_ populations, respectively. Together, *pleio4.1* and *pleio14.1* explained 22.01% of the phenotypic variation for TKW, PH, and DTF, being this the strongest effect observed among all QTL. *pleio20.1* and *pleio4.1* showed the highest additive and dominance effect, respectively.

**Fig 5.**
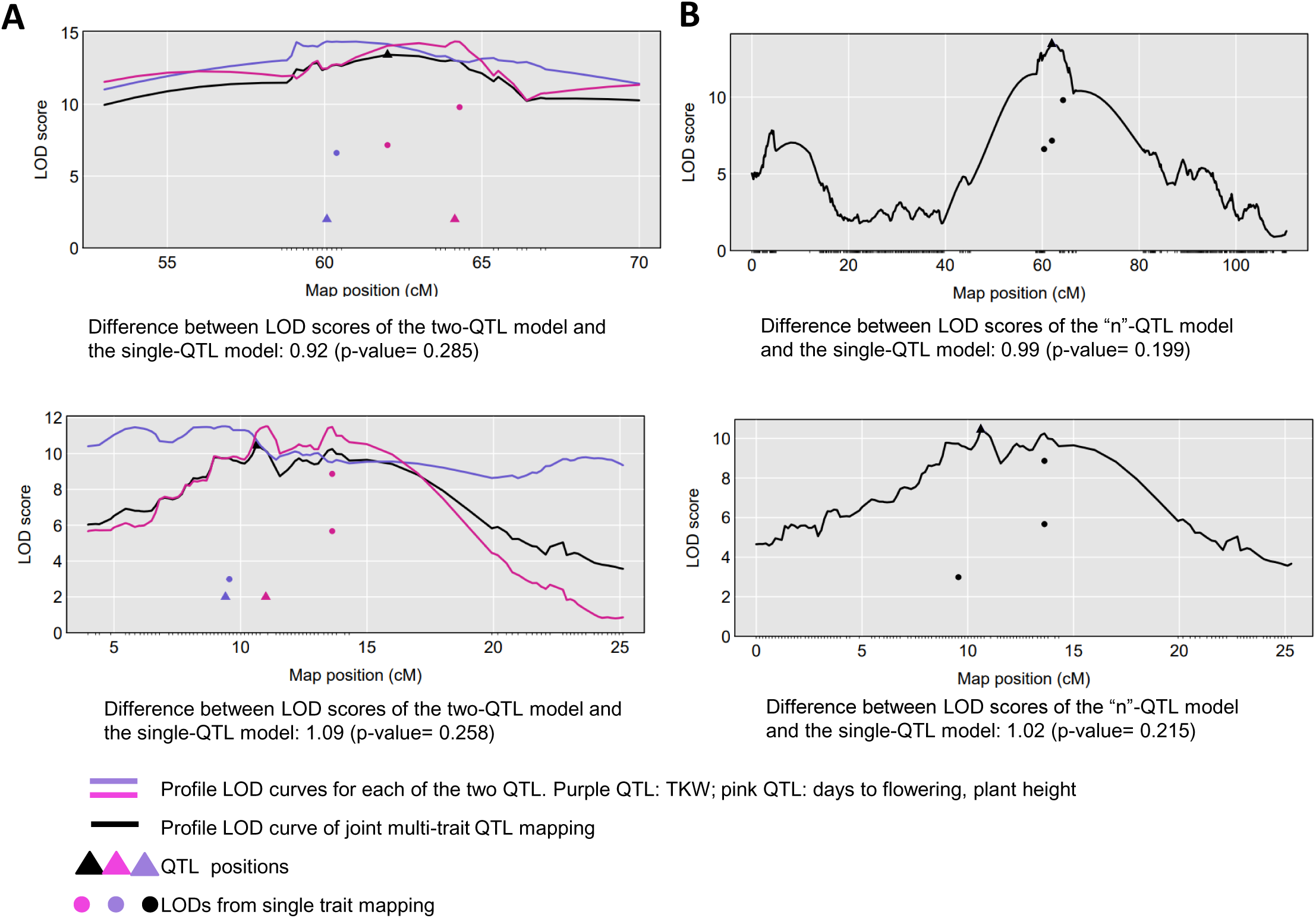
Comparative QTL analysis to detect pleiotropic QTL. Two tests were performed: (A) one vs. two QTL and (B) one vs. “n” QTL. Tests were performed considering the traits involved in the QTL found in common for F_2_ and F_3_ populations (top graphs: *pleio4.1*; bottom graphs: *pleio14.1*). The black curve is the LOD score curve for the single-QTL model, with estimated QTL location indicated by a black triangle. The blue and pink curves are profile LOD score curves for the for the two-QTL model. Dots indicate the LOD score for the traits considering a single-QTL model.

**Table 4.**
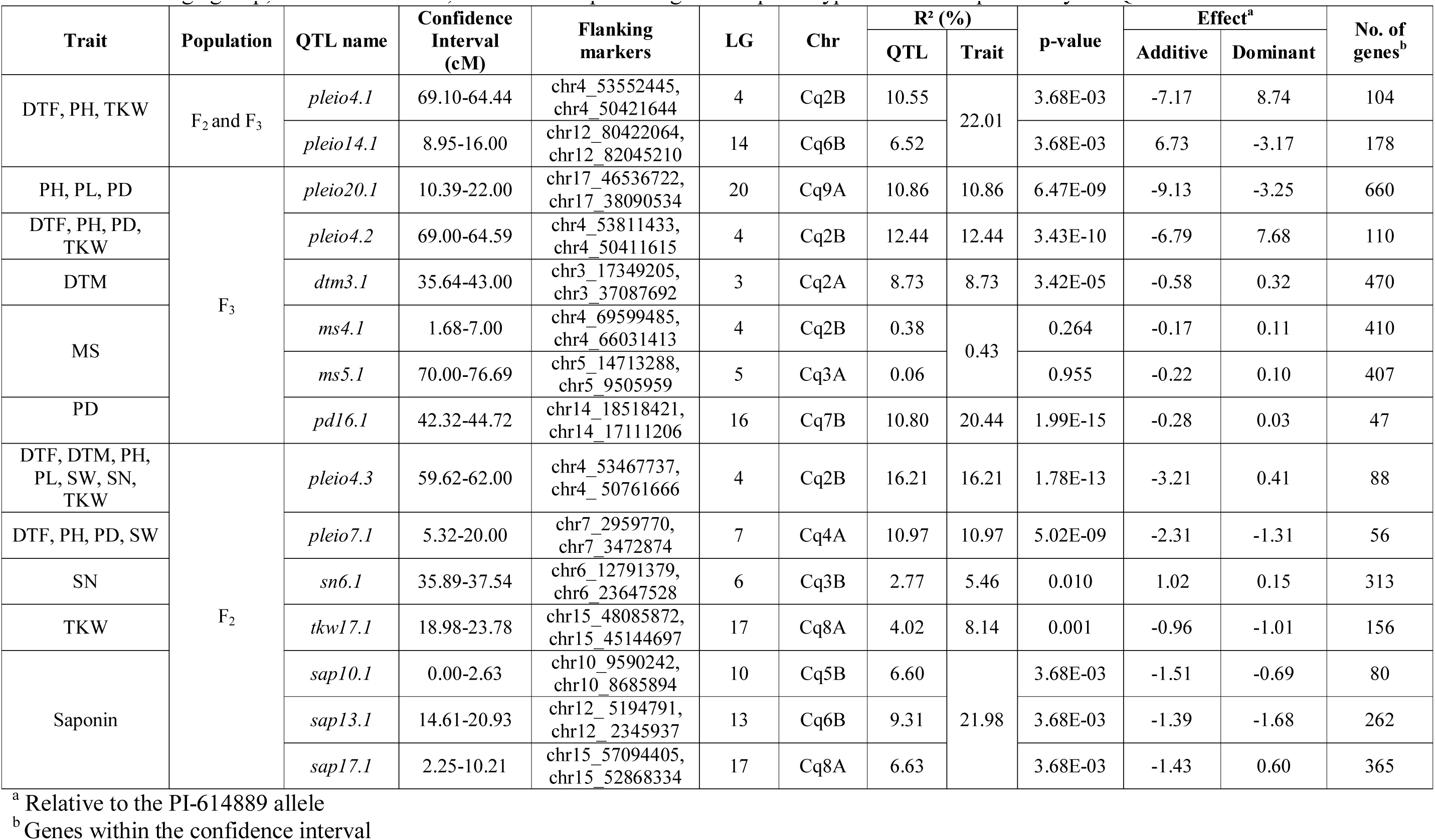
Summary statistics of QTL mapping with the F_2_ population and 328 F_3_ families. The trait acronyms are explained in the methods section. LG: linkage group; Chr: chromosome, R²: estimated percentage of the phenotypic variance explained by the QTL.

| Trait | Population | QTL name | Confidence Interval (cM) | Flanking markers | LG | Chr | R<sup>2</sup> (%) |  | p-value | Effect<sup>a</sup> |  | No. of genes<sup>b</sup> |
| --- | --- | --- | --- | --- | --- | --- | --- | --- | --- | --- | --- | --- |
|  |  |  |  |  |  |  | QTL | Trait |  | Additive | Dominant |  |
| DTF, PH, TKW | F<sub>2</sub> and F<sub>3</sub> | *pleio4.1* | 69.10-64.44 | chr4_53552445, chr4_50421644 | 4 | Cq2B | 10.55 | 22.01 | 3.68E-03 | -7.17 | 8.74 | 104 |
|  |  | *pleio14.1* | 8.95-16.00 | chr12_80422064, chr12_82045210 | 14 | Cq6B | 6.52 |  | 3.68E-03 | 6.73 | -3.17 | 178 |
| PH, PL, PD | F<sub>3</sub> | *pleio20.1* | 10.39-22.00 | chr17_46536722, chr17_38090534 | 20 | Cq9A | 10.86 | 10.86 | 6.47E-09 | -9.13 | -3.25 | 660 |
| DTF, PH, PD, TKW |  | *pleio4.2* | 69.00-64.59 | chr4_53811433, chr4_50411615 | 4 | Cq2B | 12.44 | 12.44 | 3.43E-10 | -6.79 | 7.68 | 110 |
| DTM |  | *dtm3.1* | 35.64-43.00 | chr3_17349205, chr3_37087692 | 3 | Cq2A | 8.73 | 8.73 | 3.42E-05 | -0.58 | 0.32 | 470 |
| MS |  | *ms4.1* | 1.68-7.00 | chr4_69599485, chr4_66031413 | 4 | Cq2B | 0.38 | 0.43 | 0.264 | -0.17 | 0.11 | 410 |
|  |  | *ms5.1* | 70.00-76.69 | chr5_14713288, chr5_9505959 | 5 | Cq3A | 0.06 |  | 0.955 | -0.22 | 0.10 | 407 |
| PD |  | *pd16.1* | 42.32-44.72 | chr14_18518421, chr14_17111206 | 16 | Cq7B | 10.80 | 20.44 | 1.99E-15 | -0.28 | 0.03 | 47 |
| DTF, DTM, PH, PL, SW, SN, TKW | F<sub>2</sub> | *pleio4.3* | 59.62-62.00 | chr4_53467737, chr4_50761666 | 4 | Cq2B | 16.21 | 16.21 | 1.78E-13 | -3.21 | 0.41 | 88 |
| DTF, PH, PD, SW |  | *pleio7.1* | 5.32-20.00 | chr7_2959770, chr7_3472874 | 7 | Cq4A | 10.97 | 10.97 | 5.02E-09 | -2.31 | -1.31 | 56 |
| SN |  | *sn6.1* | 35.89-37.54 | chr6_12791379, chr6_23647528 | 6 | Cq3B | 2.77 | 5.46 | 0.010 | 1.02 | 0.15 | 313 |
| TKW |  | *tkw17.1* | 18.98-23.78 | chr15_48085872, chr15_45144697 | 17 | Cq8A | 4.02 | 8.14 | 0.001 | -0.96 | -1.01 | 156 |
| Saponin |  | *sap10.1* | 0.00-2.63 | chr10_9590242, chr10_8685894 | 10 | Cq5B | 6.60 | 21.98 | 3.68E-03 | -1.51 | -0.69 | 80 |
|  |  | *sap13.1* | 14.61-20.93 | chr12_5194791, chr12_2345937 | 13 | Cq6B | 9.31 |  | 3.68E-03 | -1.39 | -1.68 | 262 |
|  |  | *sap17.1* | 2.25-10.21 | chr15_57094405, chr15_52868334 | 17 | Cq8A | 6.63 |  | 3.68E-03 | -1.43 | 0.60 | 365 |

<sup>a</sup> Relative to the PI-614889 allele
<sup>b</sup> Genes within the confidence interval

We performed a genome wide epistasis analysis to investigate how alleles influence each other in terms of enhancement and suppression and also examined how different alleles of genes influence phenotypes (DTF, PH and TKW). As the first step of this epistasis analysis, the phenotype matrix was decomposed by singular value decomposition (SVD) into eigentraits (ET). Two ET captured 69.00% and 21.00% of the total variance among DTF, PH, and TKW and were selected to perform a pairwise scan of the SNPs (Fig 6A). Then, we constructed a heatmap where 46,52% of the alleles at genome level had a minor simultaneous effect (>-1 or <1) on DTF, PH and TKW. Moreover, alleles located within the pleiotropic region *pleio4.1* showed 252 significant interactions with other alleles in all LGs, except for LGs 14, 15 and 18. Interestingly, we found that 96.82% of the alleles located within *pleio4.1* (source 4 in Fig 6B) had suppressive interaction with other alleles at the genome level (reparametrized coefficient <0). Moreover, while PI-614889 alleles at *pleio4.1* (source 4 in Fig 6B) had a negative effect on DTF and PH and a positive effect on TKW, PI-614889 alleles at *pleio14.1* (source 14 in Fig 6B) had a positive effect on DTF and PH and a negative effect on TKW. Positive and negative SNP effects refer to the reparametrized coefficient (either <0 or >0) from the pairwise regression as described by Tyler et al. (2013).

**Fig 6.**
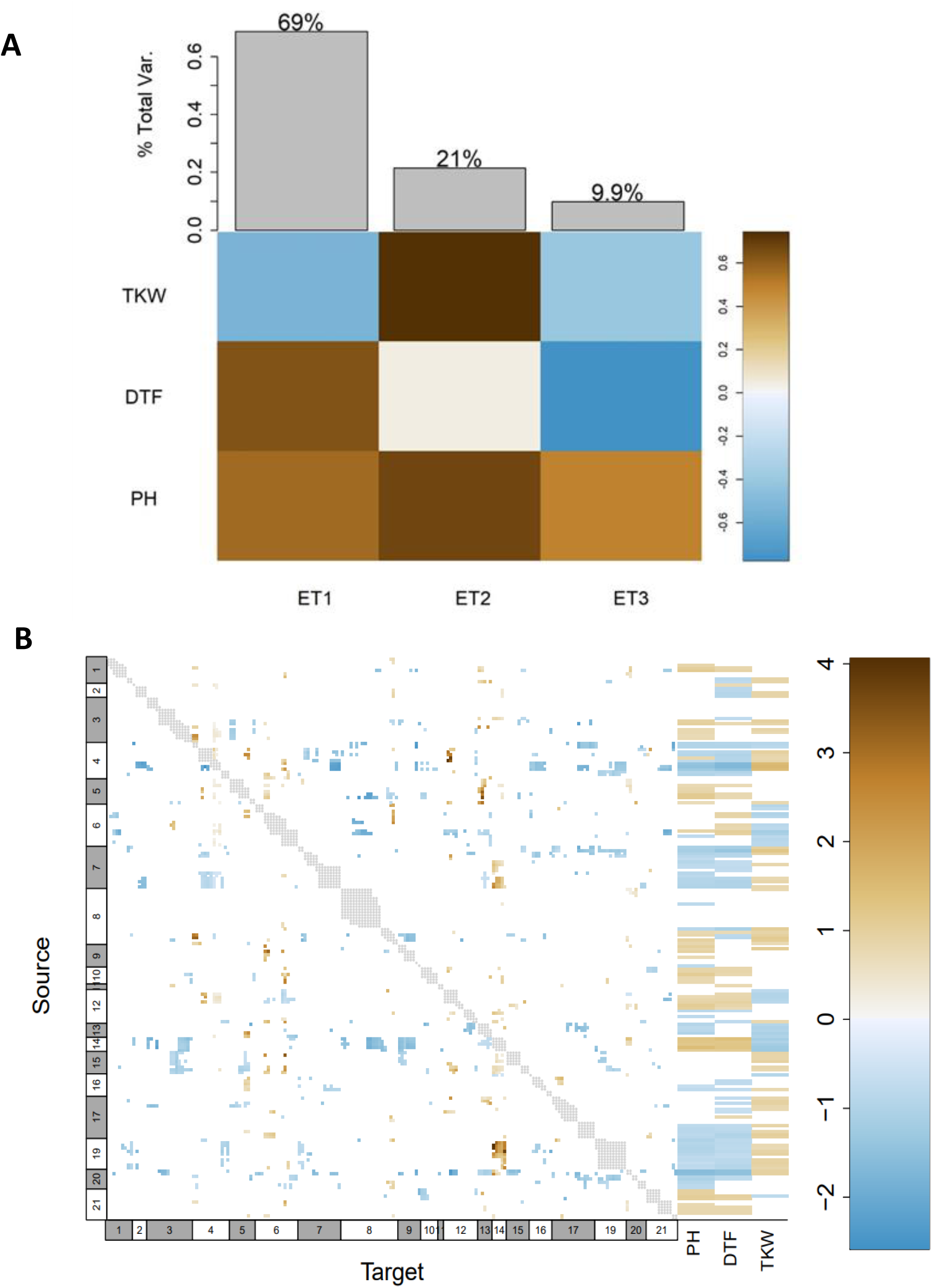
Genome wide epistasis analysis and effects of alleles at genome level on days to flowering (DTF), plant height (PH) and thousand kernel weight (TKW). (A) Decomposition of phenotypes into eigentraits (ET). Colors of the heatmap correspond to the global variance fraction of each ET. (B) Heatmap showing interactions between all pair of alleles at genome level, resulting from a pairscan analysis. Every allele was assigned as “source” and “target” for the pairscan analysis. To the right of the heatmap, interaction of every allele (assigned as source) on the phenotypes involved in the pleiotropy is shown. The heatmap scale represents the reparametrized coefficient calculated by the software and might be interpreted as the direction of the allelic effect. Grey dots show marker pairs that were not included in the pairwise scan due to complete linkage. Numbers at the x and y axes in white and grey boxes correspond to linkage groups.

### Identification of putative candidate genes controlling agronomically important traits

We searched for candidate genes within the confidence intervals of all QTL. We reasoned that trait-related SNPs could be found within or close to the genes contributing to quantitative variation (quantitative trait genes, QTG). Altogether, 1,874 genes were found within non-overlapping confidence intervals of fifteen QTL (Table S6). Nevertheless, we focused on the QTL *pleio4.1* and *pleio14.1* because of their pleiotropic effects on multiple traits and because they were common in F_2_ and F_3_ populations. Accordingly, the QTL *pleio4.1* and *pleio14.1* contributed to the phenotypic variation of three traits: DTF, PH and TKW, and 282 genes were identified within their confidence intervals.

Among the 282 genes described above, we found 41 genes with a previously described function related to flowering-time, photoperiod, and yield regulation in other plant species. Later, we compared the sequences of these genes between both parents of the population (Table S7). From all the SNPs and InDels that differed between the parents for the selected genes, we chose those that were homozygous for each parent and had a putative effect on the function of the encoded protein. From 83 selected variants, we could only identify seven SNPs in the sequencing data of the F_2_ population (Figure S12). To assess the possible effects of the variants, we grouped the F_3_ plants according to the corresponding F_2_ genotype at the variant locus and performed t-tests (α=0.05) to compare DTF, PH, and TKW between the groups. As result, none of the analyzed variants explained the phenotypic variance observed for DTF, PH and TKW (Figure S12).

Afterwards, we used available whole-genome sequencing and phenotypic data of a quinoa diversity set (310 quinoa accessions grown in a two-years experiment in Kiel, Germany) (Patirange et al. 2020) to perform the same analysis. Namely, we grouped the diversity set based on the genotypes of the F_2_ parents (either CHEN-109 or PI-614889) at the variant locus and performed t-tests (α=0.05) to compare DTF, PH, and TKW among the created groups. As result, we observed several significant phenotypic differences for PH and/or TKW and/or DTF when we grouped the quinoa diversity set based on the genotypes of our seed and pollen parents (Figure S13). Interestingly, a missense SNP variant at *TSL-KINASE INTERACTING PROTEIN 1* (*TKI1*) had simultaneous significant effects on DTF, PH and TKW (Fig 7).

**Fig 7.**
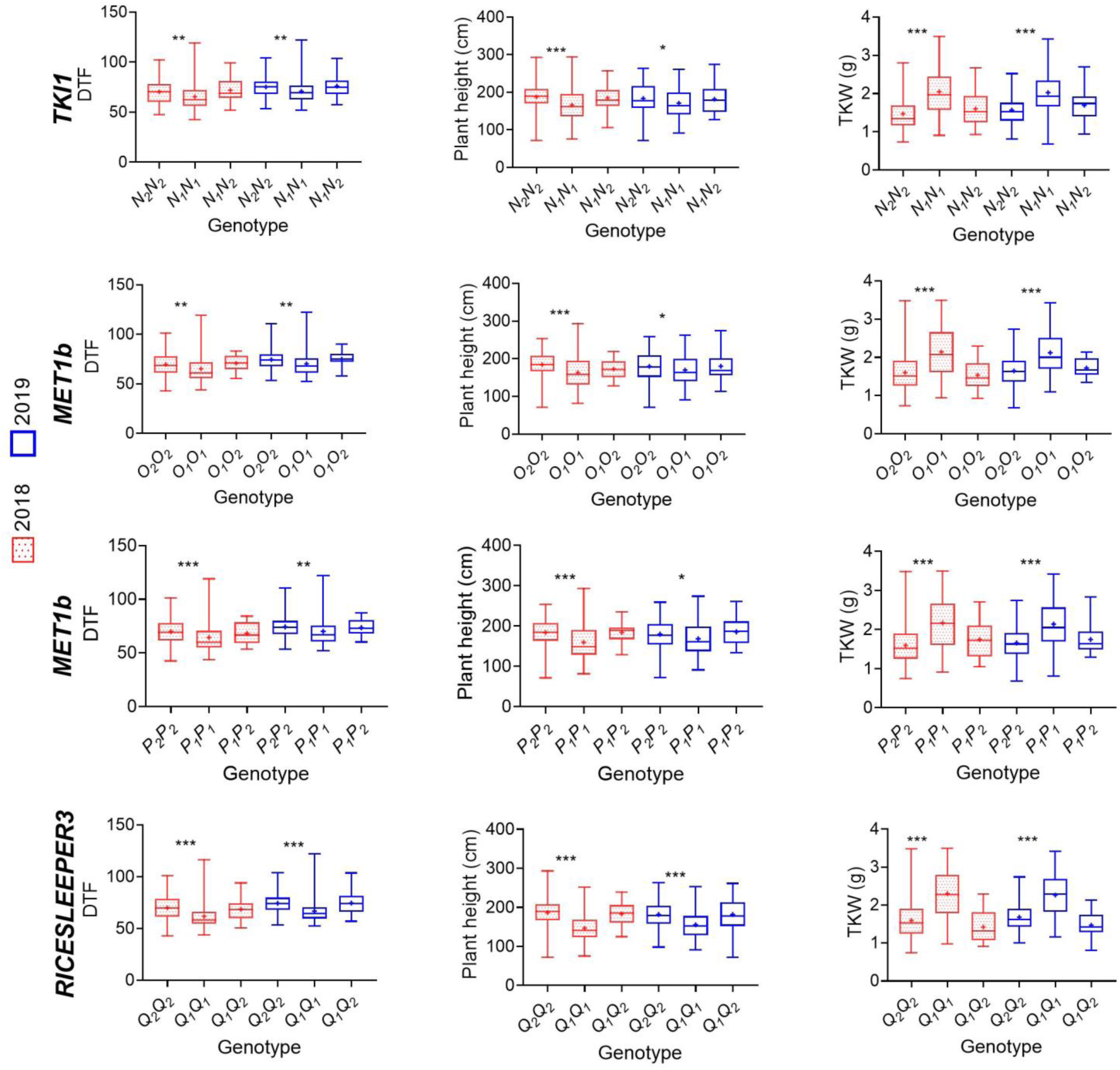
Evaluation of variant haplotypes using available whole-genome sequencing and phenotypic data of 310 quinoa accessions. Phenotypic effects of haplotype variations within four candidate genes are shown: *TSL-KINASE INTERACTING PROTEIN 1* (*TKI1*) (SNP: chr12_81633685), *DNA (CYTOSINE-5)-METHYLTRANSFERASE 1* (*MET1b*) (frameshift variant chr4_56534732), *DNA (CYTOSINE-5)-METHYLTRANSFERASE 1* (*MET1b*) (disruptive inframe deletion chr4_56534915), and *RICESLEEPER3* (disruptive_inframe_insertion chr4_55091902). The variants genotypes correspond to, for instance, *N_1_N_1_* (our homozygous parent PI-614889), *N_1_N_2_* (heterozygous), *N_2_N_2_* (our homozygous parent CHEN-109) and are described in Table S4. Significant differences between genotypes are shown by asterisks (t-test, α<0.05=***, α<0.01=**, α<0.001=***). Phenotypic data of different years are shown by different colors. DTF: days to flowering, TKW: thousand kernel weight

Accessions in the quinoa diversity set that carried the PI-614889 genotype (*N_1_N_1_*) at chr12_81633685 flowered earlier, were shorter and showed higher TKW values than those carrying the CHEN-109 (*N_2_N_2_*) genotype. Furthermore, a frameshift variant and a disruptive in-frame deletion at *DNA (CYTOSINE-5)-METHYLTRANSFERASE 1* (*MET1b*) had simultaneous significant effects on DTF, PH and TKW. Accessions in the quinoa diversity set carrying the deletion and the frameshift variant (CHEN-109 genotype) flowered later, were taller and had lower TKW values than those accessions without the deletion and the frameshift variant (PI-614889 genotype). A similar scenario was observed for a disruptive in-frame insertion at *RICESLEEPER3,* where the accessions carrying the insertion (CHEN-109 genotype) had higher values of DTF, PH and lower TKW values (Fig 7).

## Discussion

We exploited the recent advances in sequencing technologies and computational analysis methods to localize QTL for agronomically important traits in quinoa. A high-density genetic map was constructed with a segregating F_2_ population, and 15 QTL were mapped with phenotypic data from two segregating generations. Candidate genes underlying the quantitative variation were identified within the QTL.

We calculated broad-sense heritabilities and genetic advance (GA) with a selection intensity of 5%, which resulted moderate to high across all traits (excluding days to maturity). Interestingly, the high heritability coupled with high GA observed for days to flowering, panicle density and saponin content indicates that selection may be more effective for these traits. Moreover, previous studies reported heritability values for days to flowering of 70.1% (two-year experiment with five quinoa genotypes) (Al-Naggar et al. 2017) and of 91.0% (quinoa diversity set phenotyped for two-years) (Patirange et al. 2020). The same studies calculated heritabilities of 89.7% for TKW and 68.0% for panicle density. Thus, the stated values in our study are in accordance with previous reports. Nevertheless, Al-Naggar et al. (2017) and Patirange et al. (2020) reported values of 85.0% and 60.7% for plant height. Differently, in our study, the traits plant height and panicle length showed moderate heritability, both with values of around 38%. A possible explanation is that the pollen parent showed a wide range and large variability in plant height, resulting in lower estimates of heritability.

Our study considered Skim-seq as a genome complexity-reduction method for constructing a genetic map for quinoa. In our hands, genotyping by Skim-seq was effective for QTL mapping and could be applied for a minor crop like quinoa, for which available resources and commercial interest is currently limited (Böndel and Schmid 2021). We showed, by several quality controls, that the challenge of calling high-quality heterozygous SNP at low sequencing coverage (∼1.07x) could be overcome by modifications to conventional bioinformatic pipelines and imputation. Moreover, our results showed than whole-genome sequencing with a coverage as low as ∼1.0x would be sufficient for QTL mapping. QTL mapping using by whole-genome low-coverage sequencing has been successfully applied in chickpea and tobacco. In these studies, RILs and backcross populations were sequenced with depths from ca. 0.75 to 1.0x, and the number of markers for mapping ranged from ∼4,000 to ∼53,000 (Kale et al. 2015; Tong et al. 2021). Although Skim-seq was sufficient for constructing a high-density genetic map, there are limitations to this method. First, there are no tools available for the accurate imputation of InDels in F_2_ populations of polyploid crops. Thus, further uses of our genotypic data are restricted only to SNPs. The second important drawback of this approach is that the centromeric regions cannot be accurately sequenced due to their repetitive nature. However, these problems might be addressed soon by development of more sophisticated bioinformatics tools for imputation (Jordan et al. 2022).

Using Skim-seq and phenotyping, we mapped 15 QTL for ten different traits with a wide range of explained phenotypic variation (from 0.43% for MS to 22.01% for PH, DTF and TKW) and a wide range of the contribution of individual QTL to the total phenotypic variation (from 0.06% to 16.21%). QTL explaining the total phenotypic variance are unreachable because variation in quantitative traits is affected by many small-effect QTL (often undetectable) with additive and dominance effects, and QTL-QTL interactions.

Importantly, in our study, the use of families in the F_3_ generation allowed plants of the same family to be considered as replicates, allowing measurement of environmental variances and thus, significantly increasing the power and precision of QTL detection. Accordingly, the accuracy of our QTL analysis is supported by the QTL *sap10.1*, found for saponin content. This QTL, which was previously reported in other studies (Jarvis et al. 2017; Patirange et al. 2020), harbors a *TRITERPENE SAPONIN BIOSYNTHESIS ACTIVATING REGULATOR 2* (*TSAR2*) basic helix–loop–helix (bHLH) transcription factor, which likely controls the production of anti-nutritional triterpenoid saponins in quinoa seeds. However, no QTL was found for saponin content in the F_3_ generation. This might be due to the method, which was used to phenotype saponin content in this generation. We bulked seeds from ten plants in the field corresponding to one F_3_ family. From these bulks, we took samples of 20 seeds for saponin analysis. Thus, bulked samples of 20 seeds might not have been true representatives of thousands of seeds from a F_3_ family. Moreover, the relatively low correlation coefficient between the saponin content measured in F_2_ and F_3_ generations might be also result of the low efficiency of this method for measuring the saponin content in the F_3_ population. We suggest measuring the saponin content separately in multiple single plants of every F_3_ family to obtain more accurate values for this trait in F_3_.

We consider the common QTL among F_2_ and F_3_ populations, which depicted pleiotropy, as the most relevant QTL in our study. These QTL, explaining the phenotypic variation of days to flowering, plant height and TKW, were located at Cq2B and Cq6B. In contrast, a putative pleiotropic locus controlling days to flowering, days to maturity, plant height and panicle length was found at Cq2A by (Patirange et al. 2020). The different outcomes might respond to the different type of population that was used in our study as compared to (Patirange et al. 2020). Accordingly, we could expect different alleles segregating in our biallelic F_2_ population than the ones studied by (Patirange et al. 2020), who used a quinoa diversity set comprising more than 300 accessions. Furthermore, we found a strong correlation between days to flowering, plant height, and TKW (Fig 1), which were the traits implicated in pleiotropy. This reinforces their QTL co-localization at Cq2B and Cq6B. Similar correlations, where quinoa plants flowering earlier are shorter and depict higher yield related traits have been observed by Manjarres-Hernández et al. (2021) and Patirange et al. (2020).

We reported additive and dominance effects in a wide range of values, where the highest additive effect was observed for *pleio20.1* (-9.13), a QTL explaining phenotypic variation of plant height, panicle length and panicle density, and the highest dominant effect was reported for *pleio4.1* (8.74). However, these results solely show effects of single loci. Since quantitative variation in phenotypes result from highly complex networks and epistasis, we aimed to investigate the allele effects (relative to the PI-614889 allele) at genome-wide level on days to flowering, plant height, and TKW. We observed that nearly half of the studied alleles had a minor simultaneous effect on these three phenotypes; thus, confirming the nature of these traits as quantitative. From this analysis, we could also corroborate that *pleio4.1* and *pleio14.1* are major QTL, which themselves explain 10.55% and 6.52% of the phenotypic variance, respectively. Moreover, while PI-614889 alleles at *pleio4.1* had a negative effect on DTF and PH and a positive effect on TKW, PI-614889 alleles at *pleio14.1* had a positive effect on DTF and PH and a negative effect on TKW. This is correlated with the additive effects calculated for the markers with the highest LOD score within each QTL (Table 4). The observed contrasting effects of PI-614889 alleles at *pleio4.1* and *pleio14.1* would probably complicate breeding processes, whose aim is to reduce days to flowering and plant height and increase TKW, simultaneously (Murphy et al. 2018; Patiranage et al. 2021). In a broader sense, our results from the genome-wide epistasis analysis revealed the complexity of the regulation of days to flowering, plant height, and TKW in quinoa, which was expected given the intricated processes, such as DNA methylation, histone modification, and non-coding RNA-associated gene silencing, which underlie these traits (Pandey et al. 2021).

A search for candidate genes within the confidence intervals of the QTL was performed. We reasoned that trait-related SNPs could be found within the genes contributing to quantitative variation. Within the 15 different QTL described in our study, we found homologs of known flowering time (*HEADING DATE 3A, WRKY TRANSCRIPTION FACTOR 13*, *FLOWERING LOCUS D*), plant architecture (*APETALA2-1*) and yield-related (*SMALL BASIC INTRINSIC 1-2, SWEET*) genes from other species (He et al. 2003; Li et al. 2016; Ma et al. 2017; Patil et al. 2015; Wang et al. 2015; Yuan et al. 2014). Besides, we identified *FLOWERING LOCUS T* (*CqFT2A*) within *pleio7.1*, a QTL found in the F_2_ population, exclusively. Although *FT* genes are described in studies related to flowering time regulation in quinoa and in *C. rubrum*, no flowering-time function has been specified particularly for the *CqFT2A* paralog (Patiranage et al. 2021; Štorchová 2020). To continue towards identification of candidate genes, we mainly focused on the two pleiotropic QTL detected in F_2_ and F_3_ populations and identified putative candidate genes based on their known functions in flowering time and yield regulation in other species. Then, we investigated the effect of different sequence variants in these genes in a quinoa diversity set. As outcome and despite of the fact that non-related accessions could have different haplotypes although they possess the same SNP or InDel in the candidate gene, we found several sequence variants that significantly explained the phenotypic variation of PH and/or TKW and/or DTF in the quinoa diversity set. Hence, the genes containing these variants (either SNP or InDel) might be interesting candidates for further studies. Among these genes, we found that sequence variation at *TSL-KINASE INTERACTING PROTEIN 1* (*TKI1*) had significant simultaneous effects on days to flowering, plant height and TKW, which were the phenotypes whose variation was partly explained by the QTL *pleio14.1* in our study. In Arabidopsis, *TKI1* interacts with *TOUSLED* (*TSL*) and *TSL* loss of function mutations has pleiotropic effects on both leaf and flower development. Loss of *TSL* function also affects flowering time since it is required in the floral meristem for correct initiation of floral organ primordia (Ehsan et al. 2004; Roe et al. 1993). Therefore, it is tempting to speculate that *TKI1* is involved in the regulation of the flowering time, flower development and consequently, seed set in quinoa. Moreover, we found that sequence variations at *MET1b* and *RICESLEEPER3* have similar simultaneous effects on days to flowering, plant height and TKW, as observed for the variant at *TKI1*. In Arabidopsis, *MET1* homozygous mutants displayed late-flowering phenotypes caused by ectopic expression of *FLOWERING WAGENINGEN* (*FWA)*, a regulator of flowering time. Hypomethylation, which correlates with the mentioned late-flowering phenotypes, is often accompanied by other developmental alterations (Kakutani et al. 1996; Saze et al. 2003; Soppe et al. 2000). Furthermore, *DAYSLEEPER. DAYSLEEPER,* the Arabidopsis homolog of *RICESLEEPER3,* encodes for a transposase-like protein essential for plant growth and development. Moreover, loss of function mutants of *DAYSLEEPER* showed retarded growth and delayed flowering (Bundock and Hooykaas 2005; Knip et al. 2012). Importantly, the evidence of the role of these genes in the regulation of different biological processes is given for the model plant Arabidopsis while the observed pleiotropic regulation of days to flowering, plant height and TKW in our study might be controlled by quinoa-specific genes. Hence, if *TKI1, MET1* and *RICESLEEPER3* have the same function in quinoa can only be verified by further investigations. A first step towards elucidating the molecular mechanism governed by these genes might be expression analysis, for instance. Furthermore, haplotype analyses may focus on the up- and downstream regulatory regions of the most relevant candidate genes in our study. Besides, despite of the lack of reliable transformation protocols for quinoa, screening of mutants and assessing their phenotypic effects seems to be another feasible approach for determining the function of these genes. Moreover, a recent study offers perspectives for functional studies in quinoa using virus-induced gene silencing (VIGS) (Ogata et al. 2021).

On the other hand, molecular markers linked to the pleiotropic QTL identified in the current study can be directly used in quinoa breeding programs for simultaneous selection of different traits. Moreover, the provided information about QTL effects could guide breeders towards the selection of early, short and high yielding quinoa genotypes. Future work may address fine mapping of the interesting pleiotropic regions and characterization of candidate genes. Overall, the results presented in this study will help provide a framework for future research on the molecular mechanisms of flowering and other agronomically important traits and facilitate marker assisted selection (MAS) in quinoa breeding programs.

## Acknowledgements

We thank Monika Bruisch, Brigitte Neidhardt-Olf and Elisabeth Kokai for technical assistance. Mireia Vidal-Villarejo provided support in the bioinformatic analysis of sequencing data. Prof. Dr. Mark Tester and Dr. Elodie Rey for providing the reference genome QQ74_V2 and the parental sequences, and for their continuous advice.

## Statements and Declarations

### Funding

The financial support of this work was provided by the Schleswig-Holstein Stiftung; Proj. 2019/59.

#### Author contributions

C.J and N.E directed the project and conceived the research. N.M.T designed the experiments and conducted the greenhouse experiment. N.M.T and F.B. conducted the field experiment and performed QTL mapping. K.S. carried out DNA sequencing and provided advice on subsequent bioinformatic analyses. N.M.T carried out bioinformatic analyses and constructed the genetic map. N.M.T and F.B. carried out DNA isolation and marker analysis. N.M.T, together with all authors, wrote and finalized the manuscript.

#### Data Availability

The datasets generated during and/or analyzed during the current study are available from the corresponding author on reasonable request.

## Supplementary material

**Figure S1.** Flow charts depicting the (A) bioinformatics pipeline and (B) the genetic map construction pipeline. Main steps of the pipelines are shown in solid-line boxes, method/parameters are shown in dashed boxes and the number of markers is shown in red dashed boxes. PE= pair end.

**Figure S2.** Agarose gel electrophoresis of PCR products from the InDel marker JASS5. (A) Homozygous F_3_ plants. From lane 1 to 10, plants are homozygous for allele *R_1_* (parent CHEN-109; 189 bp); and from lane 11 to 20, plants are homozygous for allele *R_2_* (parent PI-614889; 164 bp). (B) Plants from segregating F_3_ families. L = middle range DNA ladder, P1 = parent CHEN-109, P2 = parent PI-614889, C = negative control. Agarose gels were run for 60 min at 100 V.

**Figure S3.** Phenotypic variation in the F_2_ population. (A) Variation of panicle density. Numbers in yellow represent the scoring scale used for phenotyping. (B) Variation of plant height illustrated by seven F_2_ individuals (A to G) and the parental lines. (C) Variation in days to maturity illustrated by the different colors of the panicle. PI-614889 had reached maturity stage and was ready to harvest. CHEN-109 had not reach seed filling stage. Individual C is at seed filling stage. All pictures were taken 16 weeks after sowing.

**Figure S4.** SNP densities across 18 quinoa chromosomes. The number of SNP within 1 Mb window size are shown by different colors. Densities were calculated by CMplot R package using the raw data (∼17 million SNP).

**Figure S5.** Percentage of genome-wide missing data. Black dots show missing markers. Black horizontal lines represent several continuous missing markers. Chromosomes are separated by black vertical lines. Numbers above correspond to the quinoa chromosomes (even numbers: Cq1A to Cq9A; uneven numbers: Cq1B to Cq9B).

**Figure S6.** Data filtering before map construction. (A) Number of markers per individual. (B) Number of genotyped individuals for each marker. (C) Histogram of the proportion of markers for which pairs of individuals have matching genotypes. Dashed red lines show the filtering thresholds. Data below the dashed red line in A and B, and to the right of the dashed line in C was removed.

**Figure S7.** Quinoa linkage map based on 334 plants from an F_2_ population derived from a cross between CHEN-109 and PI-614889. The map consists of 133,913 markers arranged in 5,219 bins and it was drawn with LinkageMapView R package. Numbers above indicate linkage groups (LG) and chromosome numbers. Horizontal blue lines show the location of the first marker of each bin followed by the number of markers in each bin in parenthesis. Marker names are coded as “S” + Chromosome number + “_” + physical position of the marker. Genetic distances in cM are written to the left of each LG.

**Figure S8.** Map density across 21 linkage groups from the F_2_ population derived from a cross between CHEN-109 and PI-614889. Densities were recorded by LinkageMapView R package. Binned-markers are shown by horizontal black lines.

**Figure S9.** Quality control of the marker data used for linkage map construction. (A) Marker’s segregation distortion and missing proportion (number of individuals that are missing a marker in a specific locus / 334 individuals) calculated by ASMap R package; different colors correspond to each linkage group. (B) Estimated number of crossovers and double crossovers calculated by ASMap R package; blue dots represent the estimated values for each F_2_ individual.

**Figure S10.** Frequencies of the parental alleles calculated from the F_2_ population using the program ABHgenotype R package in each of the linkage groups. Different alleles are shown by different colors. Linkage group numbers are shown to the right. The x axis shows physical positions of the 133,913 markers according to the reference genome QQ74_V2. Hetero: heterozygous genotype.

**Figure S11.** Comparative QTL analysis to detect pleiotropy for (A) *pleio20.1*, (B) *pleio4.2*, (C) *pleio4.3* and (D) *pleio7.1*. Two tests were performed: one vs. two QTL (to the left) and one vs. “n” QTL (to the right). The black curve is the LOD score curve for the single-QTL model, with estimated QTL location indicated by a black triangle. The blue and pink curves are profile LOD score curves for the for the two-QTL model. Dots indicate the LOD score for the traits considering a single-QTL model. DTF: days to flowering, DTM: days to maturity, PH: plant height, PL: panicle length, PD: panicle density, SN: seed number per plant, SW: seed weight per plant, TKW: thousand kernel weight.

**Figure S12.** Evaluation of variant haplotypes using the sequences of 328 F_2_ individuals and the corresponding phenotypic data of 328 F_3_ families. Phenotypic effects of haplotype variations within several candidate genes are shown: *DEXH-BOX ATP-DEPENDENT RNA HELICASE DEXH10* (*HEN2), SUPPRESSOR OF FRI 4 (SUF4), RCC1 DOMAIN-CONTAINING PROTEIN 3* (*RUG3)*, *WRKY TRANSCRIPTION FACTOR 13* (*WRKY13)* and *TSL-KINASE INTERACTING PROTEIN 1 (TKI1).* The variants genotypes correspond to, for instance, *A_1_A_1_* (our homozygous parent PI-614889), *A_1_A_2_*(heterozygous), *A_2_A_2_* (our homozygous parent CHEN-109) and are described in Table S4. Significant differences between genotypes are shown by asterisks (t-test, α<0.05=***, α<0.01=**, α<0.001=***). DTF: days to flowering, TKW: thousand kernel weight, PH: plant height.

**Figure S13.** Evaluation of variant haplotypes using available whole-genome sequencing and phenotypic data of 310 quinoa accessions. Phenotypic effects of haplotype variations within several candidate genes are shown: *ATHB-15/CORONA, DEXH-BOX ATP-DEPENDENT RNA HELICASE DEXH10* (*HEN2), NRT1/ PTR 2.6* (*NPF2.6), RCC1 DOMAIN-CONTAINING PROTEIN 3* (*RUG3)*, *WRKY TRANSCRIPTION FACTOR 13* (*WRKY13), ETHYLENE-RESPONSIVE TRANSCRIPTION FACTOR 113* (*ERF113*) and *FLOWERING LOCUS D* (*FLD).* The variants genotypes correspond to, for instance, *H_1_H_1_* (our homozygous parent PI-614889), *H_1_H_2_*(heterozygous), *H_2_H_2_* (our homozygous parent CHEN-109) and are described in Table S4. Significant differences between genotypes are shown by asterisks (t-test, α<0.05=***, α<0.01=**, α<0.001=***). Phenotypic data of different years are shown by different colors. DTF: days to flowering, TKW: thousand kernel weight.

**Table S1.** Plant material used in this study. DTF: days to flowering, DTM: days to maturity, PH: plant height, PL: panicle length, PD: panicle density, TKW: thousand kernel weight, SW: seed weight per plant, SN: seed number per plant, SC: Saponin content, MS: Mildew susceptibility.

**Table S2.** Methods for phenotypic evaluation.

**Table S3.** Polymerase chain reaction (PCR) and agarose gel electrophoresis description.

**Table S4.** Allele and genotype nomenclature used in this study.

**Table S5.** Genetic and phenotypic segregation for two traits in the F_2_ and F_3_ populations. Red axil pigmentation was determined five weeks after sowing. The InDel marker JAASS5 was described by Zhang et al. (2017). *R_1_* and *R_2_* represent the 189 bp and 164 bp alleles, respectively.

**Table S6.** Complete list of genes located within the non-overlapping QTL confidence intervals.

**Table S7.** Genes within *pleio4.1* and *pleio14.1* confidence intervals with a previous described function related to flowering time, photoperiod response or yield regulation and their gene variants. Gene variants between CHEN-109 (pollen parent) and PI-614889 (seed parent) and to the reference genome QQ74_V2 are shown.

