## Supplementary Figures for "High-density mapping of QTL controlling agronomically important traits in quinoa (*Chenopodium quinoa* Willd.)": Supplementary_Figures.pdf

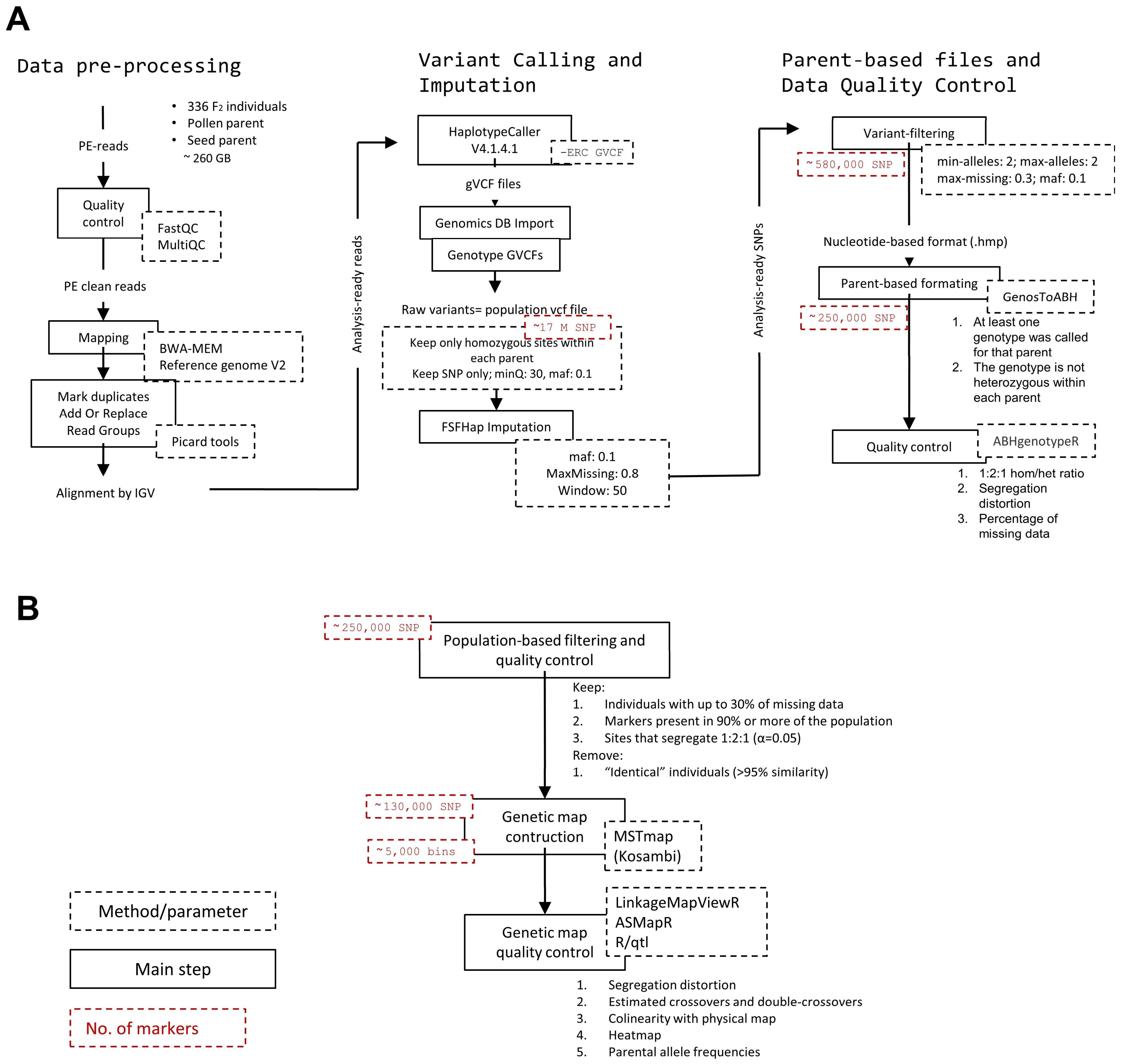

**Figure S1** Flow charts depicting the (A) bioinformatics pipeline and (B) the genetic map construction pipeline. Main steps of the pipelines are shown in solid-line boxes, method/parameters are shown in dashed boxes and the number of markers is shown in red dashed boxes. PE= pair end.

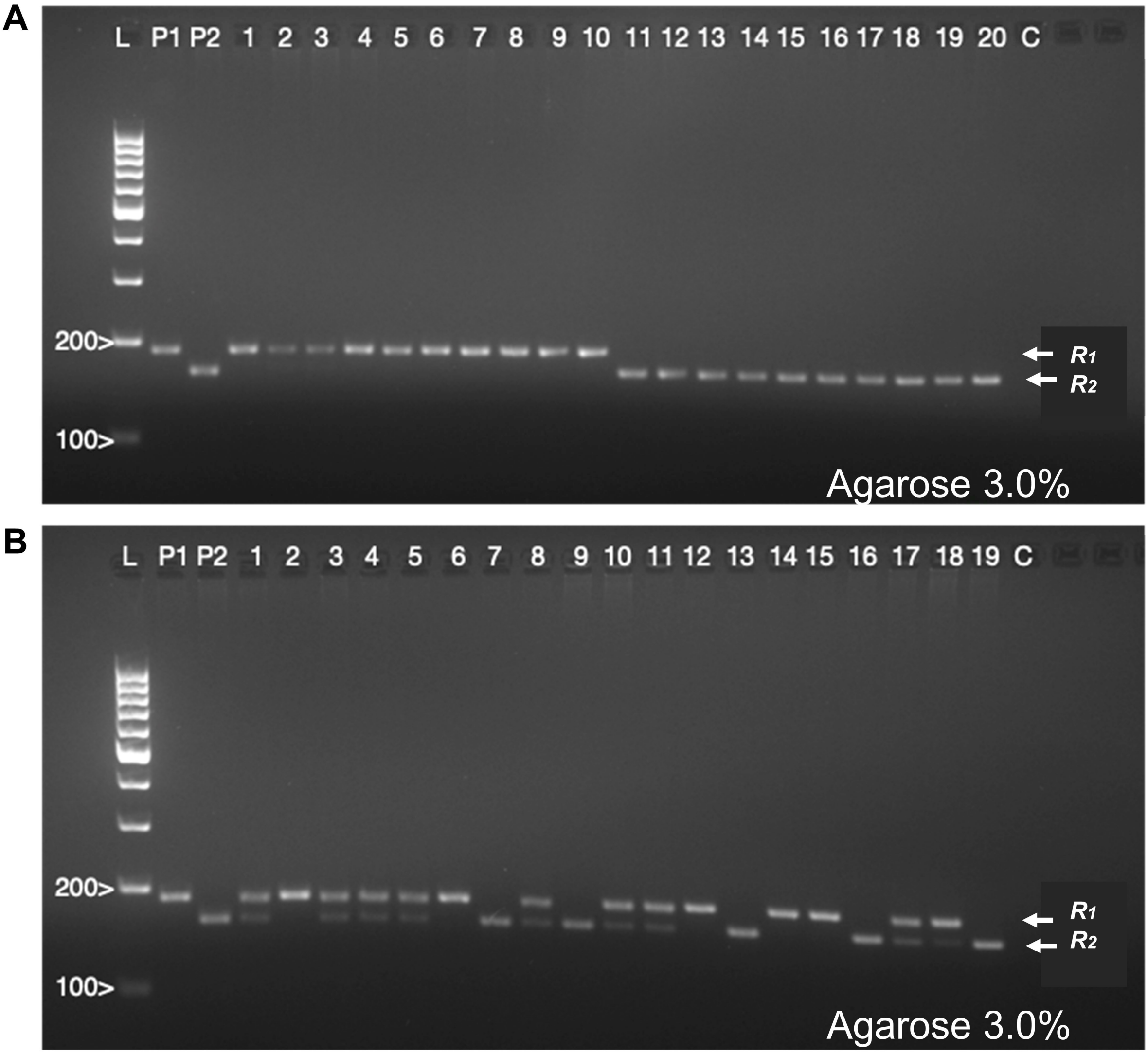

**Figure S2** Agarose gel electrophoresis of PCR products from the InDel marker JASS5. (A) Homozygous $F_3$ plants. From lane 1 to 10, plants are homozygous for allele $R_1$ (parent CHEN-109; 189 bp); and from lane 11 to 20, plants are homozygous for allele $R_2$ (parent PI-614889; 164 bp). (B) Plants from segregating $F_3$ families. L = middle range DNA ladder, P1 = parent CHEN-109, P2 = parent PI-614889, C = negative control. Agarose gels were run for 60 min at 100 V.

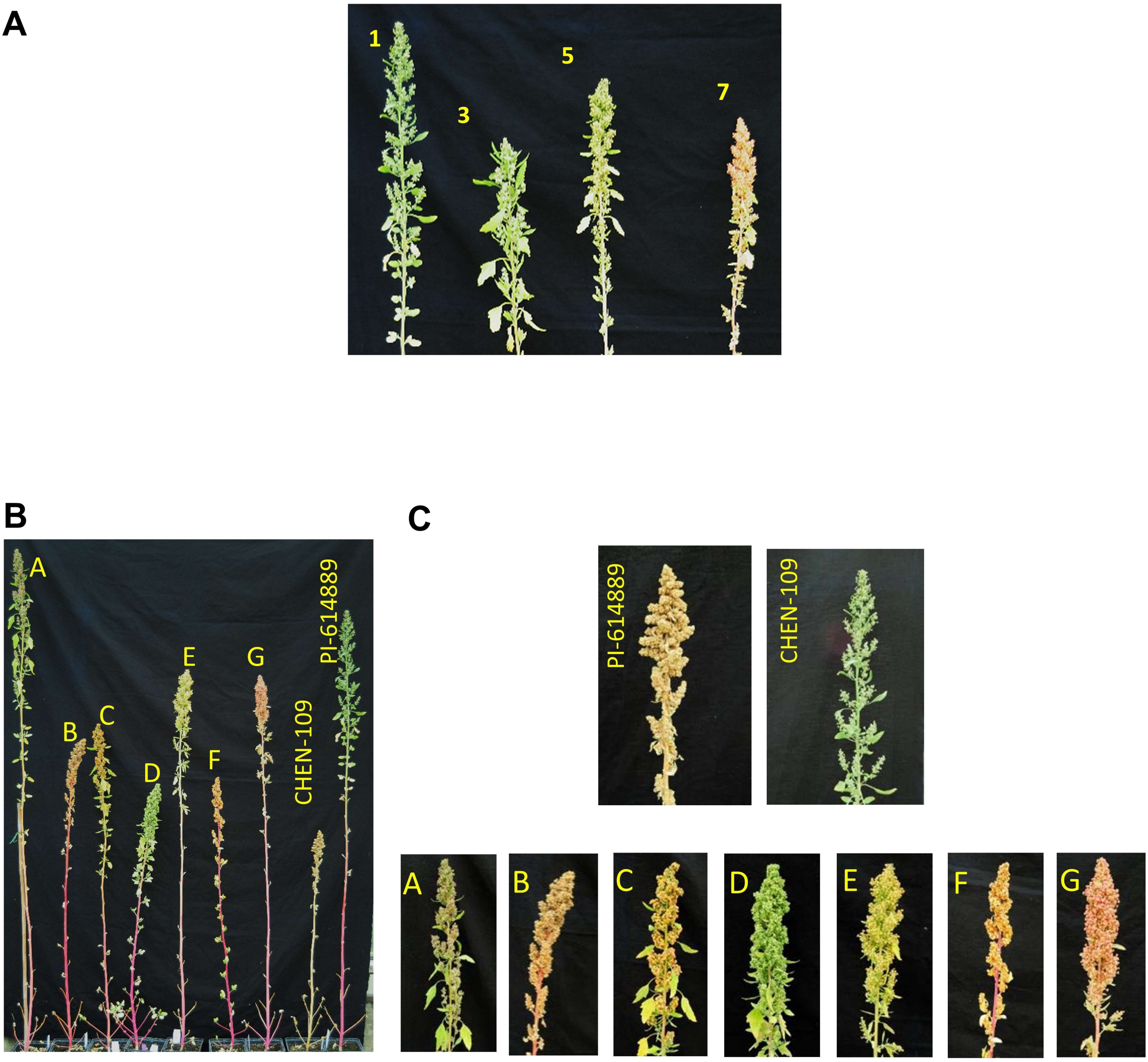

**Figure S3** Phenotypic variation in the $F_2$ population. (A) Variation of panicle density. Numbers in yellow represent the scoring scale used for phenotyping. (B) Variation of plant height illustrated by seven $F_2$ individuals (A to G) and the parental lines. (C) Variation in days to maturity illustrated by the different colors of the panicle. PI-614889 had reached maturity stage and was ready to harvest. CHEN-109 had not reach seed filling stage. Individual C is at seed filling stage. All pictures were taken 16 weeks after sowing.

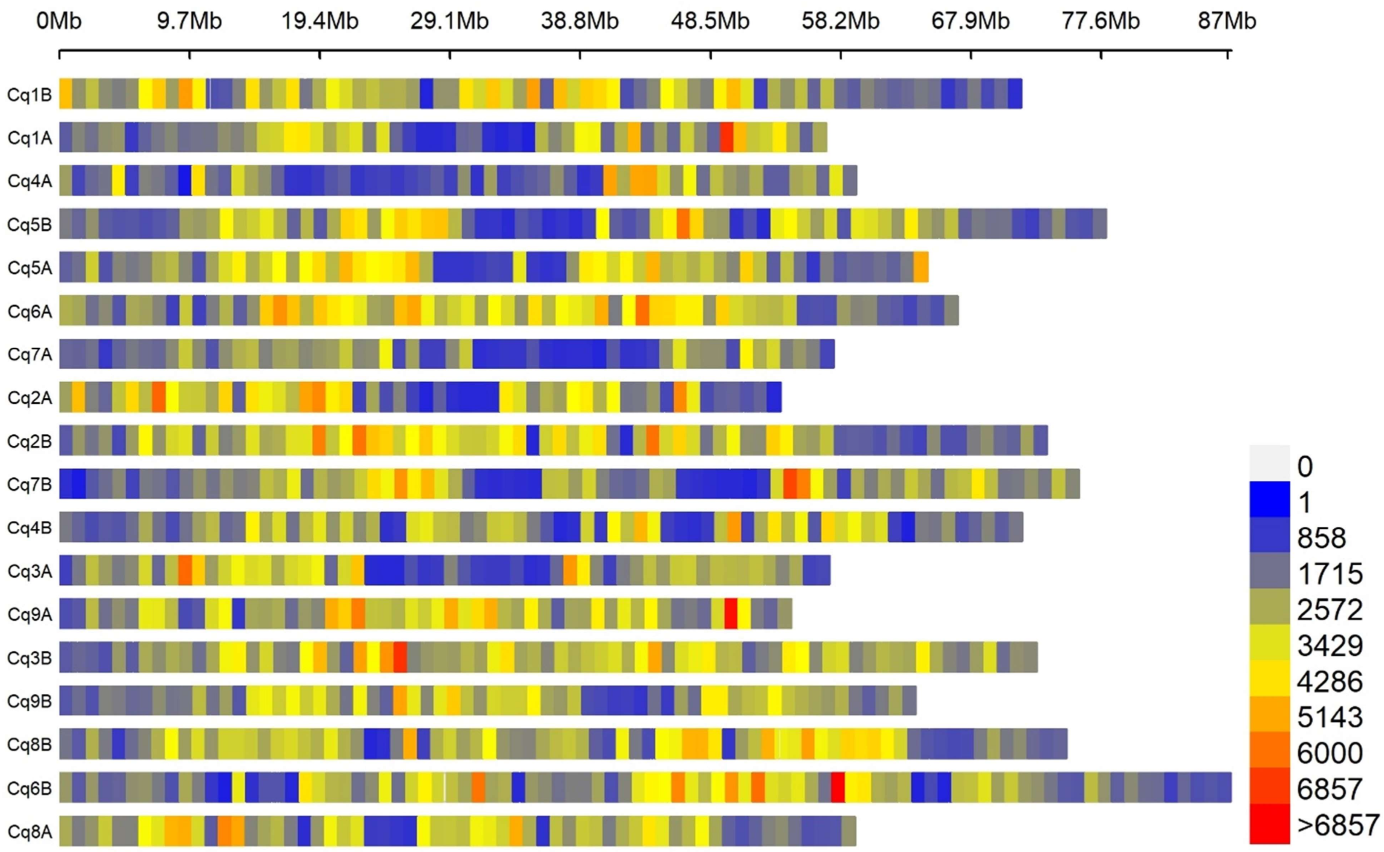

**Figure S4** SNP densities across 18 quinoa chromosomes. The number of SNP within 1 Mb window size are shown by different colors. Densities were calculated by CMplot R package using the raw data (~17 million SNP).

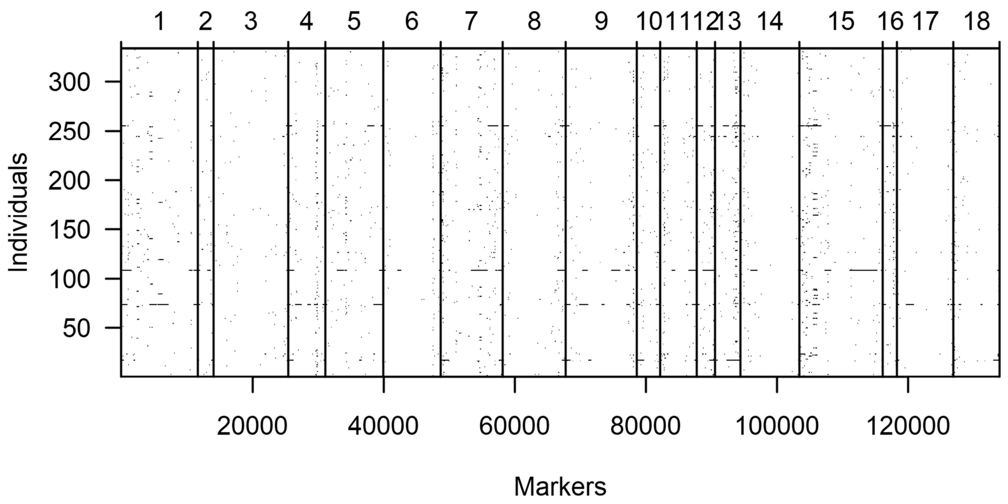

**Figure S5** Percentage of genome-wide missing data. Black dots show missing markers. Black horizontal lines represent several continuous missing markers. Chromosomes are separated by black vertical lines. Numbers above correspond to the quinoa chromosomes (even numbers: Cq1A to Cq9A; uneven numbers: Cq1B to Cq9B).

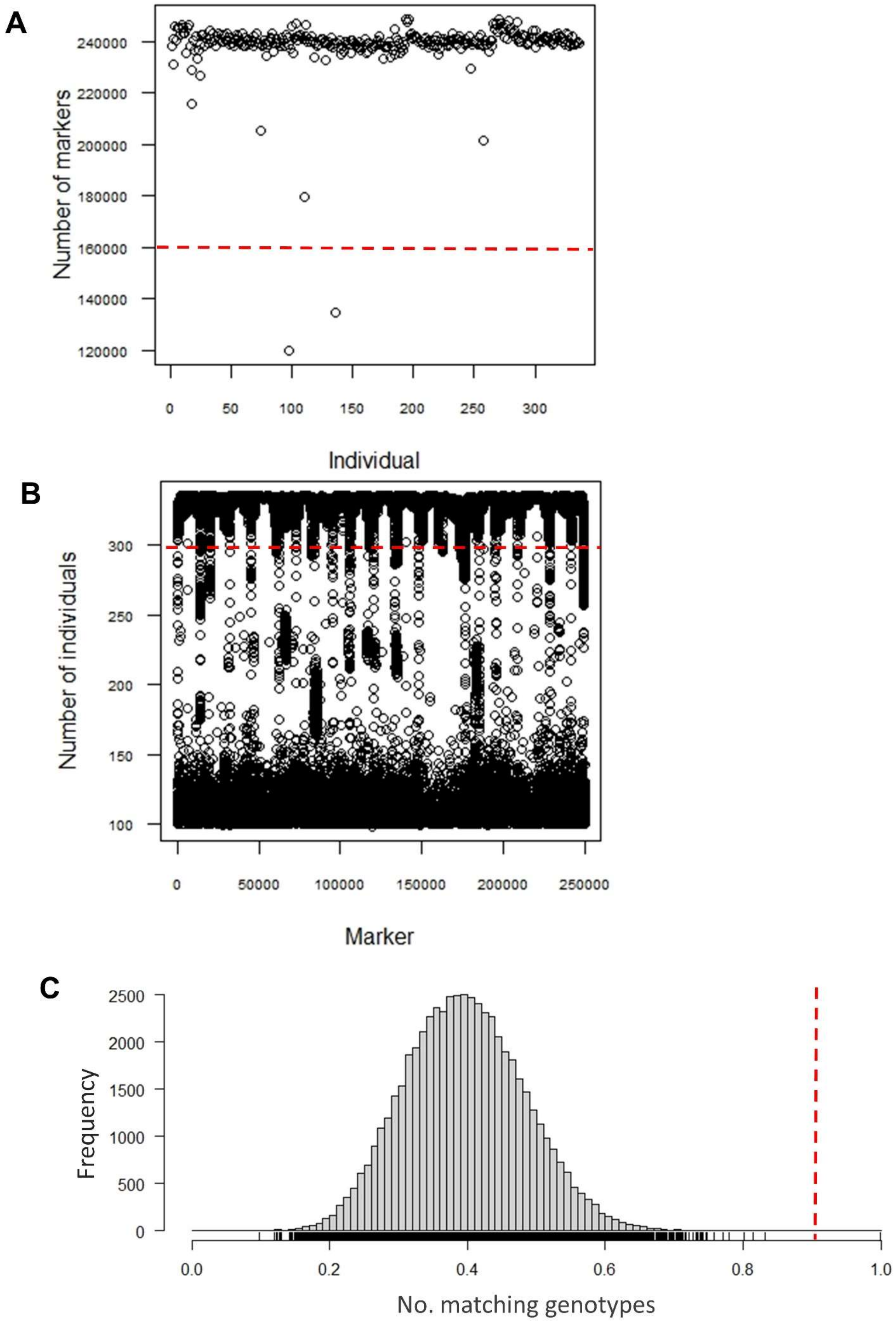

**Figure S6** Data filtering before map construction. (A) Number of markers per individual. (B) Number of genotyped individuals for each marker. (C) Histogram of the proportion of markers for which pairs of individuals have matching genotypes. Dashed red lines show the filtering thresholds. Data below the dashed red line in A and B, and to the right of the dashed line in C was removed.

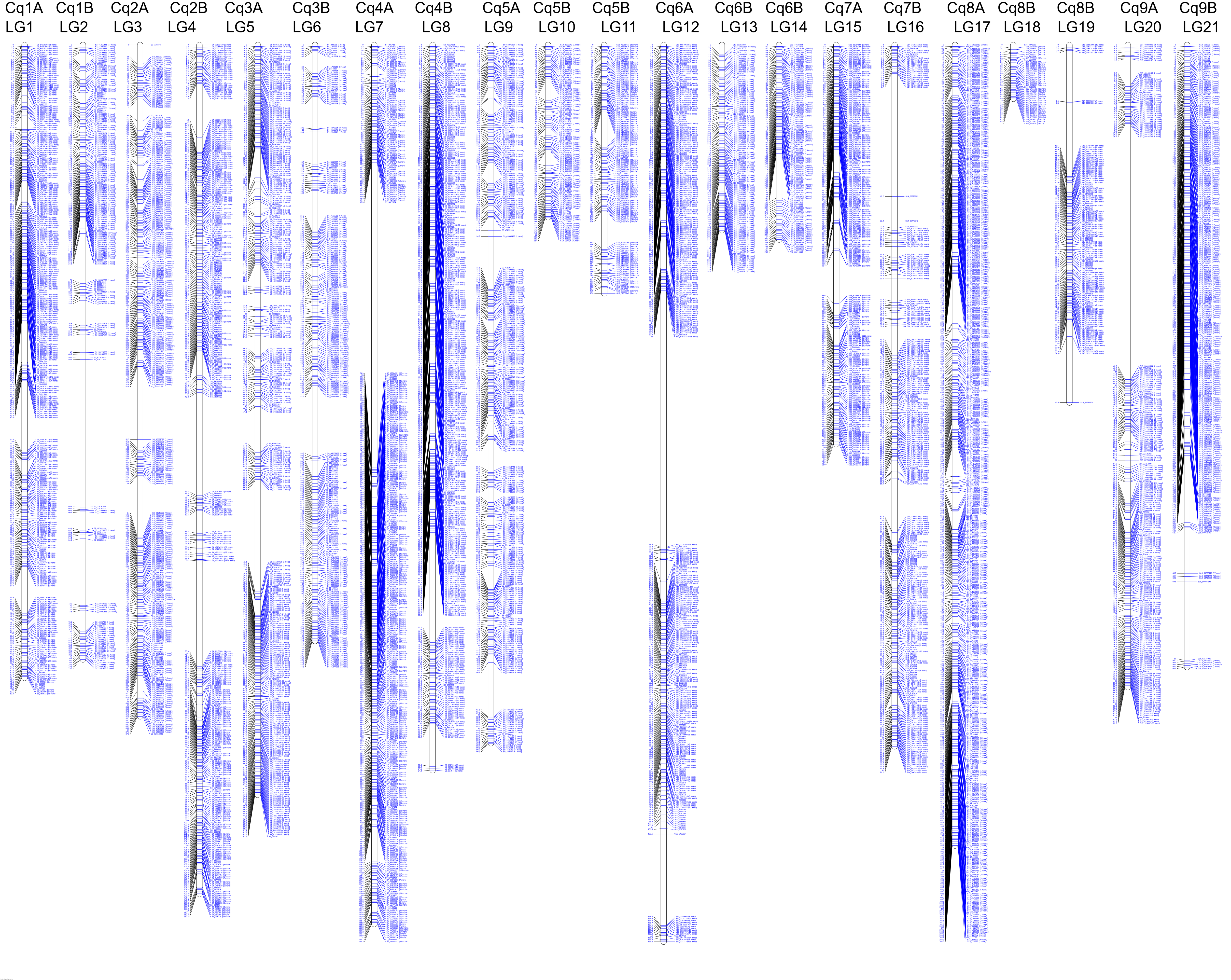
Cq1A LG1
Cq1B LG2
Cq2A LG3
Cq2B LG4
Cq3A LG5
Cq3B LG6
Cq4A LG7
Cq4B LG8
Cq5A LG9
Cq5B LG10
Cq5B LG11
Cq6A LG12
Cq6B LG13
Cq6B LG14
Cq7A LG15
Cq7B LG16
Cq8A LG17
Cq8B LG18
Cq8B LG19
Cq9A LG20
Cq9B LG21

**Figure S7** Quinoa linkage map based on 334 plants from an $F_2$ population derived from a cross between CHEN-109 and PI-614889. The map consists of 133,913 markers arranged in 5,219 bins and it was drawn with LinkageMapView R package. Numbers above indicate linkage groups (LGs) and chromosome numbers. Horizontal blue lines show the location of the first marker of each bin followed by the number of markers in each bin in parenthesis. Marker names are coded as "S" + Chromosome number + "_" + physical position of the marker. Genetic distances in cM are written to the left of each LG.

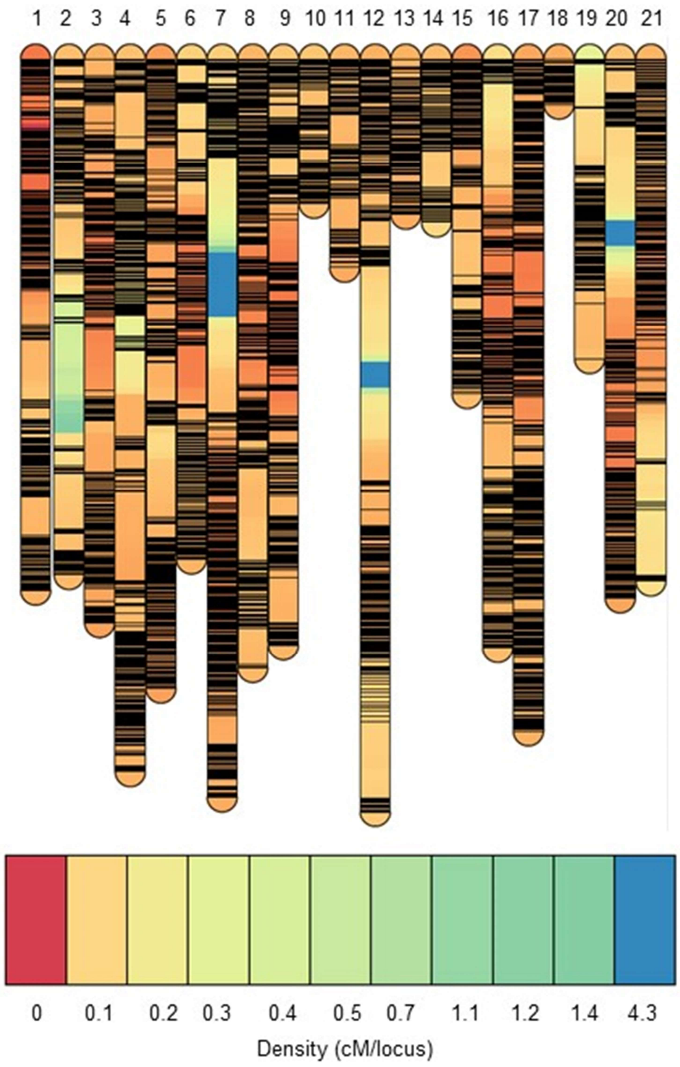

**Figure S8** Map density across 21 linkage groups from the $F_2$ population derived from a cross between CHEN-109 and PI-614889. Densities were recorded by LinkageMapView R package. Binned-markers are shown by horizontal black lines.

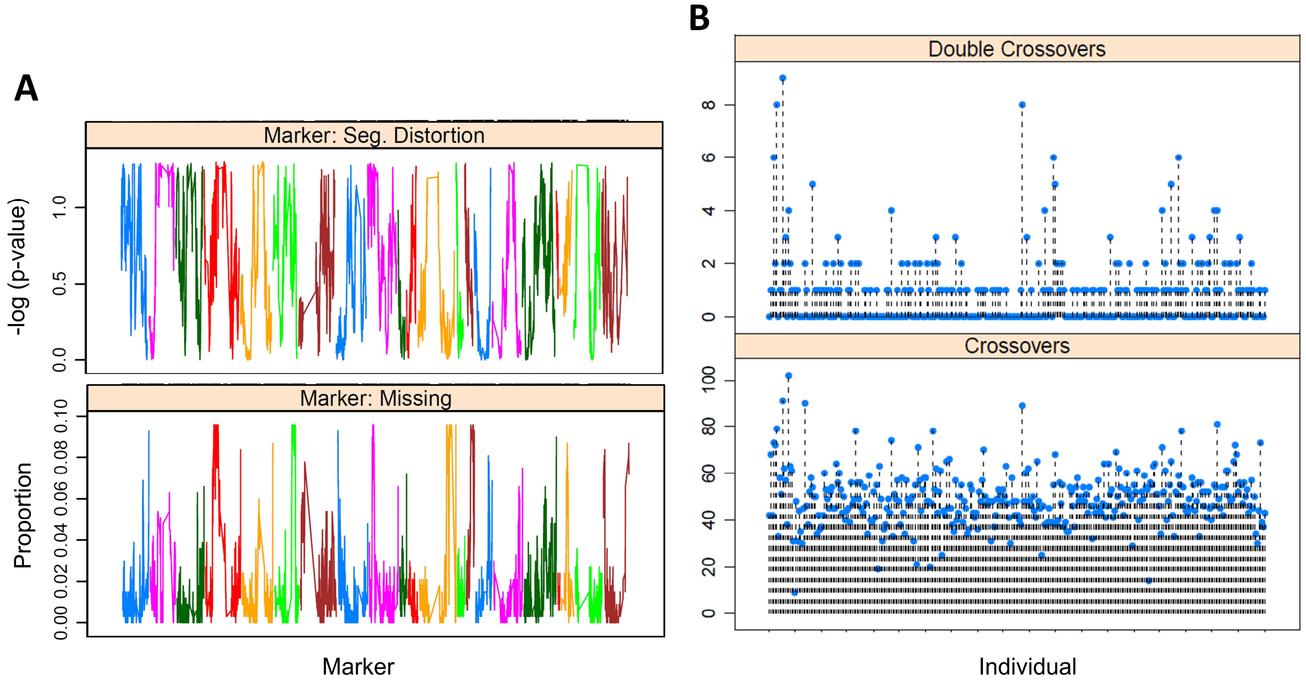

**Figure S9** Quality control of the marker data used for linkage map construction. (A) Marker's segregation distortion and missing proportion (number of individuals that are missing a marker in a specific locus / 334 individuals) calculated by ASMap R package; different colors correspond to each linkage group. (B) Estimated number of crossovers and double crossovers calculated by ASMap R package; blue dots represent the estimated values for each $F_2$ individual.

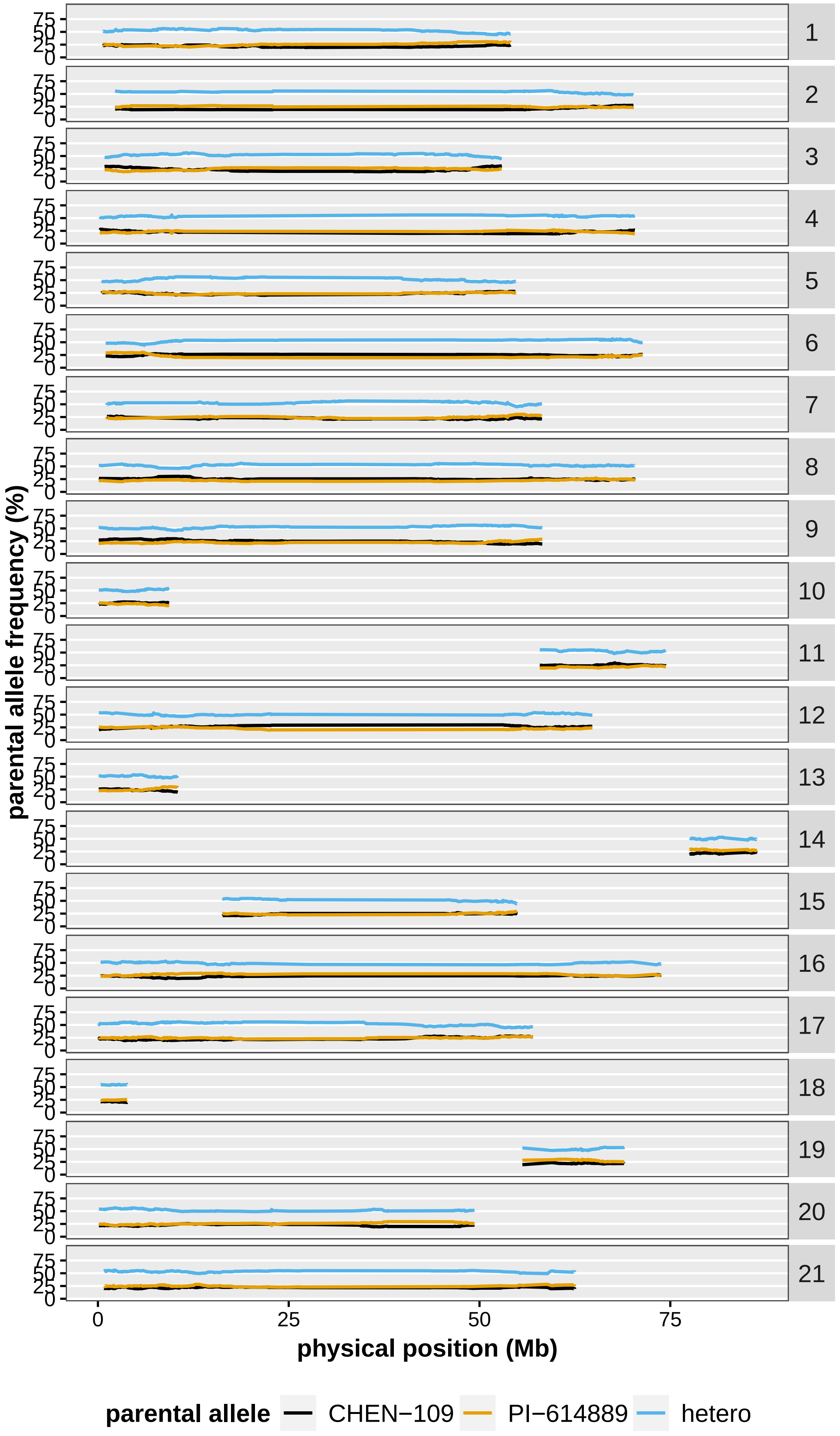

parental allele frequency (%)
0
25
50
75
physical position (Mb)
parental allele
CHEN−109
PI−614889
hetero

**Figure S10** Frequencies of the parental alleles calculated from the $F_2$ population using the program ABHgenotype R package in each of the linkage groups. Different alleles are shown by different colors. Linkage group numbers are shown to the right. The x axis shows physical positions of the 133,913 markers according to the reference genome QQ74_V2. Hetero: heterozygous genotype.

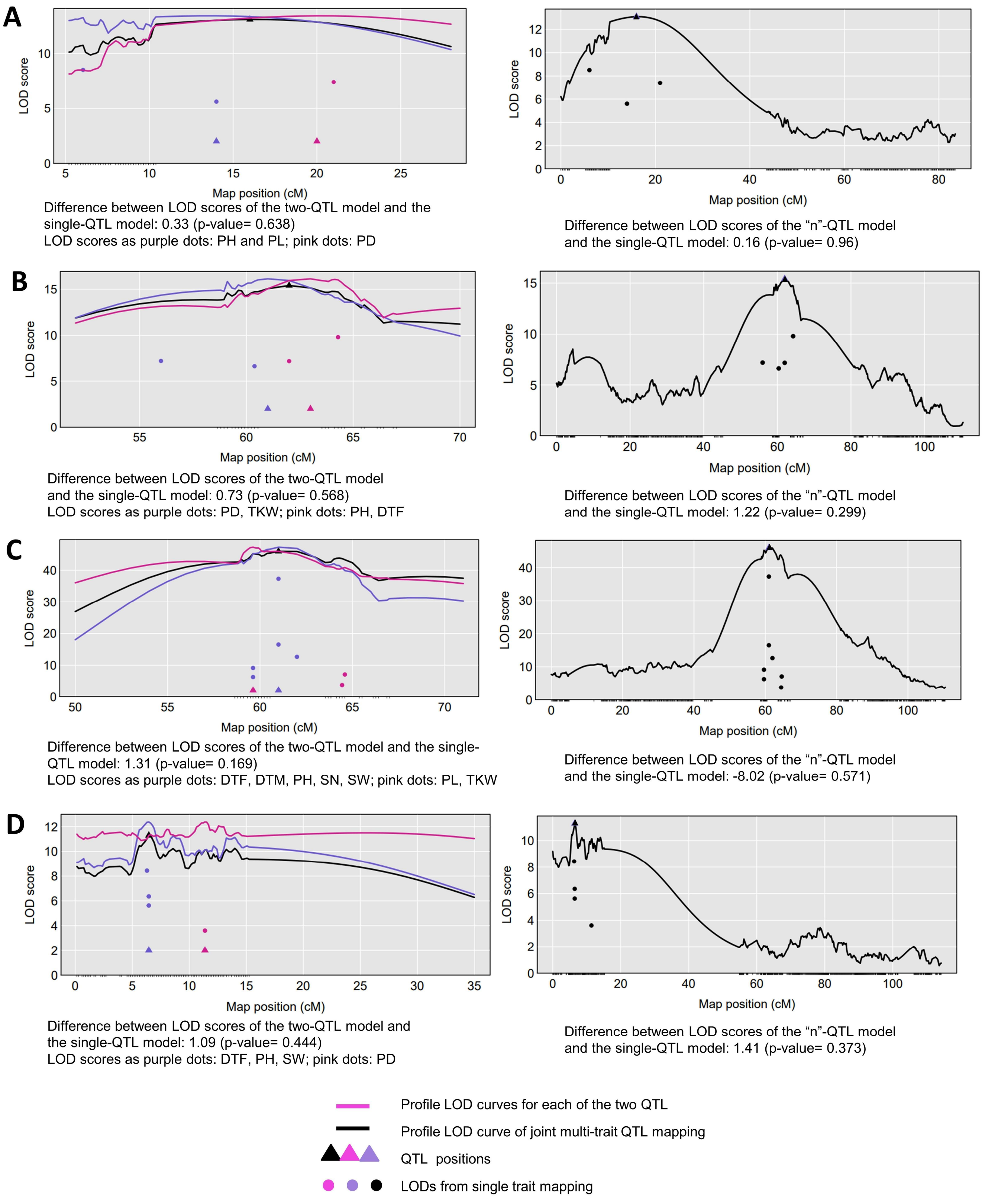

**Figure S11** Comparative QTL analysis to detect pleiotropy for (A) *pleio20.1*, (B) *pleio4.2*, (C) *pleio4.3* and (D) *pleio7.1*. Two tests were performed: one vs. two QTL (to the left) and one vs. “n” QTL (to the right). The black curve is the LOD score curve for the single-QTL model, with estimated QTL location indicated by a black triangle. The blue and pink curves are profile LOD score curves for the for the two-QTL model. Dots indicate the LOD score for the traits considering a single-QTL model. DTF: days to flowering, DTM: days to maturity, PH: plant height, PL: panicle length, PD: panicle density, SN: seed number per plant, SW: seed weight per plant, TKW: thousand kernel weight.

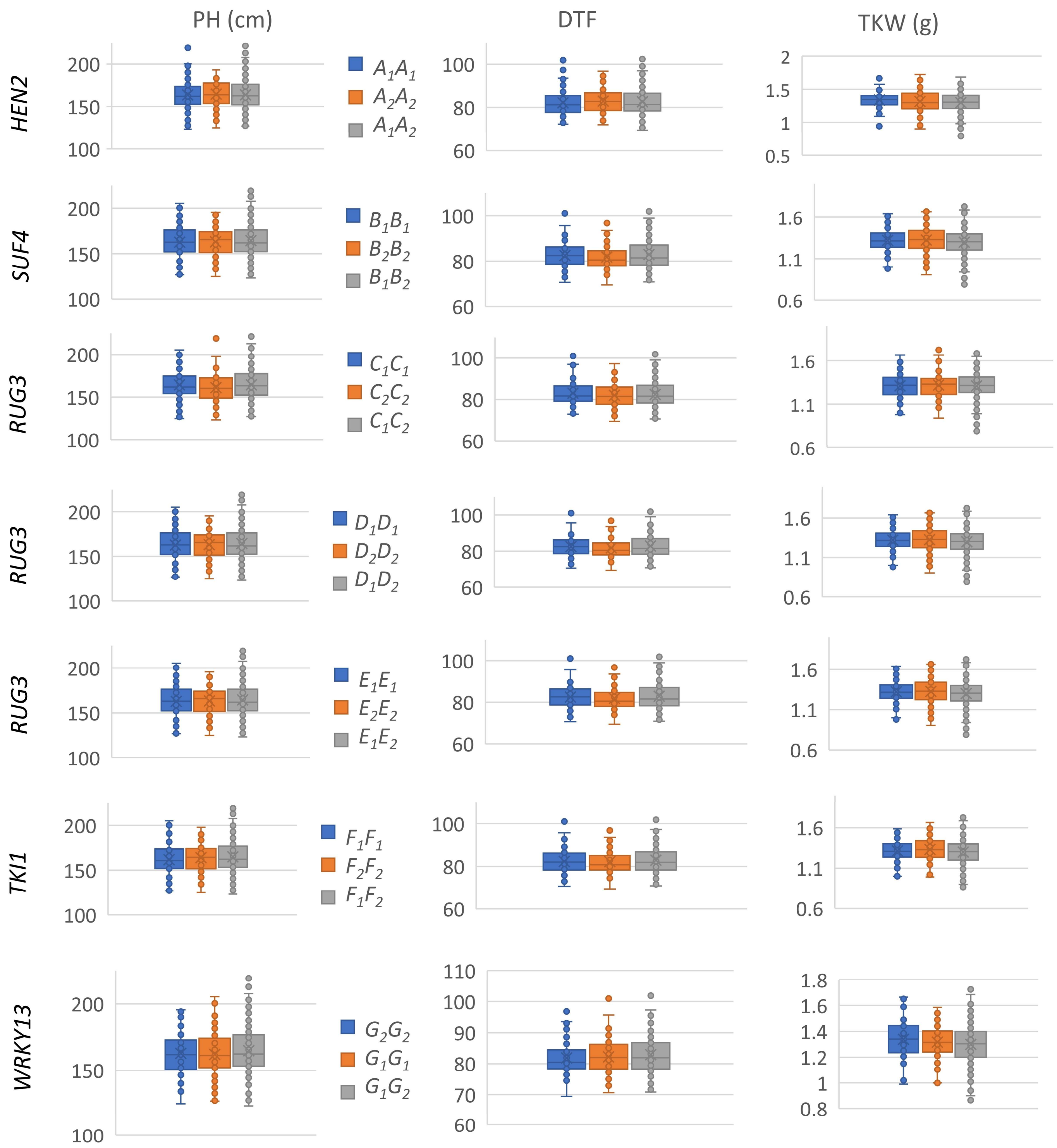

**Figure S12** Evaluation of variant haplotypes using the sequences of 328 $F_2$ individuals and the corresponding phenotypic data of 328 $F_3$ families. Phenotypic effects of haplotype variations within several candidate genes are shown: *DEXH-BOX ATP-DEPENDENT RNA HELICASE DEXH10* (*HEN2), SUPPRESSOR OF FRI 4 (SUF4), RCC1 DOMAIN-CONTAINING PROTEIN 3* (*RUG3*), *WRKY TRANSCRIPTION FACTOR 13* (*WRKY13)* and *TSL-KINASE INTERACTING PROTEIN 1 (TKI1).* The variants genotypes correspond to, for instance, $A_1A_1$ (our homozygous parent PI-614889), $A_1A_2$ (heterozygous), $A_2A_2$ (our homozygous parent CHEN-109) and are described in Supplementary Table 4. Significant differences between genotypes are shown by asterisks (t-test, α<0.05=***, α<0.01=**, α<0.001=***). DTF: days to flowering, TKW: thousand kernel weight, PH: plant height.

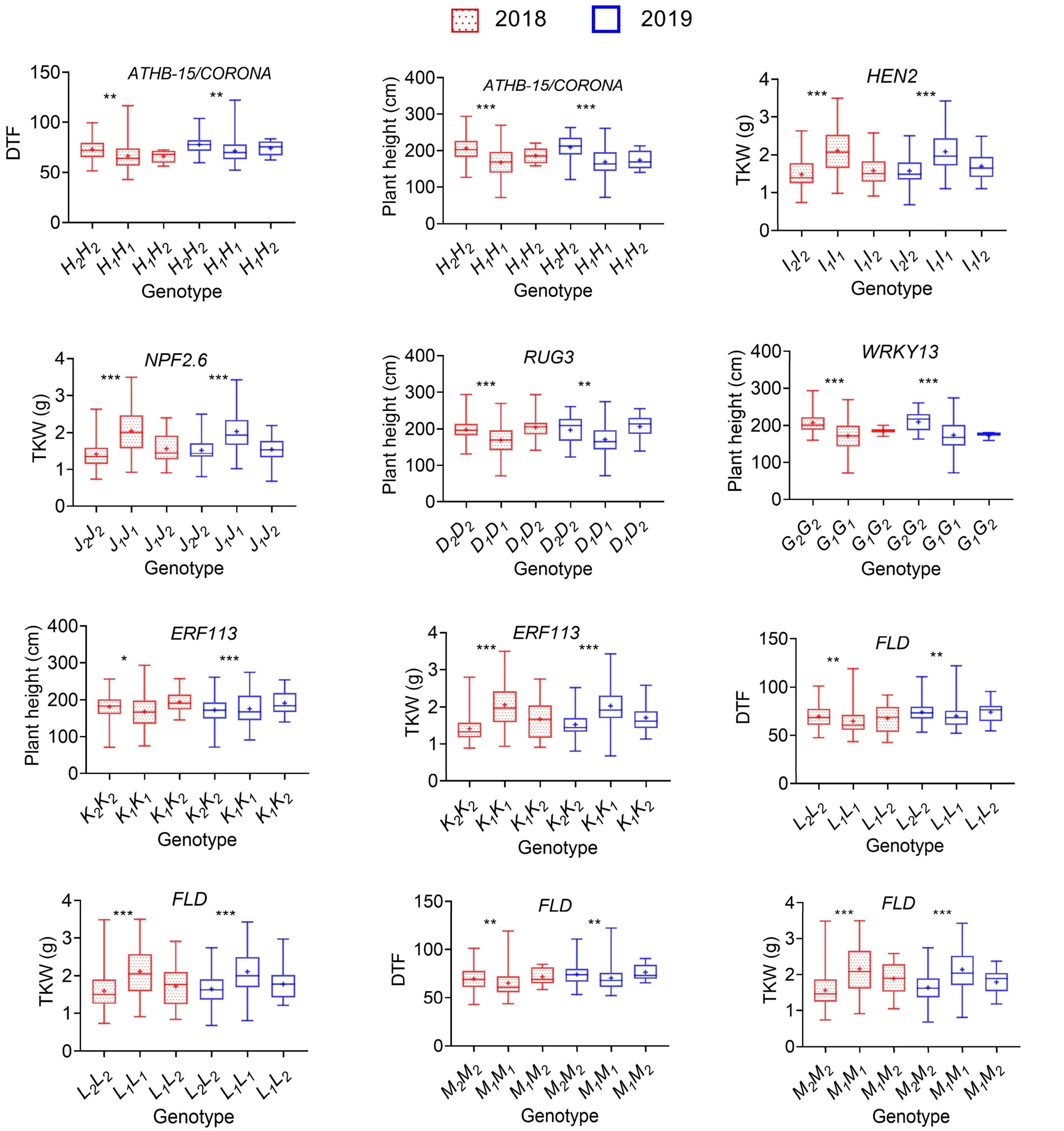

**Figure S13** Evaluation of variant haplotypes using available whole-genome sequencing and phenotypic data of 310 quinoa accessions. Phenotypic effects of haplotype variations within several candidate genes are shown: *ATHB-15/CORONA, DEXH-BOX ATP-DEPENDENT RNA HELICASE DEXH10* (*HEN2*)*, NRT1/ PTR 2.6* (*NPF2.6*)*, RCC1 DOMAIN-CONTAINING PROTEIN 3* (*RUG3*)*, WRKY TRANSCRIPTION FACTOR 13* (*WRKY13*)*, ETHYLENE-RESPONSIVE TRANSCRIPTION FACTOR 113* (*ERF113*) and *FLOWERING LOCUS D* (*FLD*)*.* The variants genotypes correspond to, for instance, $H_1H_1$ (our homozygous parent PI-614889), $H_1H_2$ (heterozygous), $H_2H_2$ (our homozygous parent CHEN-109) and are described in Supplementary Table 4. Significant differences between genotypes are shown by asterisks (t-test, α<0.05=***, α<0.01=**, α<0.001=***). Phenotypic data of different years are shown by different colors. DTF: days to flowering, TKW: thousand kernel weight.
