## Supplementary Table 1-5 for "High-density mapping of QTL controlling agronomically important traits in quinoa (*Chenopodium quinoa* Willd.)": Supplementary_Tables_1_to_5.pdf

**Table S1.** Plant material used in this study. DTF: days to flowering, DTM: days to maturity, PH: plant height, PL: panicle length, PD: panicle density, TKW: thousand kernel weight, SW: seed weight per plant, SN: seed number per plant, SC: Saponin content, MS: Mildew susceptibility.

| Seed code | Generation | No. of plants/families | No. of plants genotyped | No. of plants phenotyped | |
|---|---|---|---|---|---|
| | | | | Greenhouse | Field |
| 171115 | PI-614889 (seed parent) | - | - | 10 | 20 |
| 170876 | CHEN-109 (pollen parent) | - | - | 10 | 20 (DTM, SW and SN were not recorded) |
| 190031 | $F_2$ | 336 | 336 | 336 except for MS | - |
| 191203-191562 | $F_3$ | 334 | - | - | DTF: 5,891<br>DTM: 2,343<br>PH: 5,860<br>PL: 5,860<br>PD: 5,860<br>MS: 6,346<br>SC: 330 families were bulked together (out of 334 sown families, four were lost due to biotic stress after germination)<br>TKW: 330 families were bulked together (out of 334 sown families, four were lost due to biotic stress after germination)<br>SW: 0<br>SN: 0 |

**Table S2.** Methods for phenotypic evaluation.

| Trait | Acronym | Description |
|---|---|---|
| Days to flowering | DTF | Number of days, from sowing until the first flower opens. Recorded thrice a week in the $F_2$ and twice a week in the $F_3$ population. |
| Days to maturity | DTM | Number of days, from sowing until the panicle is completely brown. Recorded twice a week. |
| Plant height | PH | Distance in cm from root collar to panicle apex, recorded once at BBCH-81 to BBCH-89. |
| Panicle length | PL | Distance in cm from panicle base to tip, recorded once at BBCH-81 to BBCH-89. |
| Panicle density | PD | Visual score based on the observed range of panicle density recorded at BBCH-81 to BBCH-89. The score ranges from 1 to 5: (1) "Low": loose panicle with low number of spaced glomerules and panicle axes clearly visible; (3) "Intermediate": high number of glomerules tightly arranged with panicle axis often visible; (5) "High": high number of glomerules tightly packed and scarcely seen panicle axes; and (7) "Very high": very high number of glomerules compactly arranged, panicle axes not visible. Recorded once at BBCH-81 to BBCH-89. An example of panicle density scoring in the $F_2$ population can be observed in Supplementary Figure 3. |
| Mildew susceptibility | MS | Performed only in the $F_3$ population (incidence only in the field). Performed as visual score ranging from 1 to 3: (1) "High": symptoms observed in the whole plant or at least up to the upper third of the plant, (2) "Intermediate": symptoms observed in the lowest two-thirds of the plant, (3) "Low": no visible symptoms or only present in the lower third of the plant. Recorded once at BBCH-81 to BBCH-89. |
| Saponin content | SC | Recorded after harvest by foam test. Twenty seeds were placed in a 2 ml epi, shaken for 1 min at 1100 g in a GenoGrinder and incubated at room temperature for 5 min; later, the height of the formed foam (cm) was recorded (Jarvis et al., 2017). |
| Seed weight per plant | SW | Weight of the total harvest (whole panicle) per plant. Recorded once at BBCH-99. |
| Number of seeds per plant | SN | Number of seeds contained in the whole panicle per plant. Recorded once at BBCH-99. |
| Thousand Kernel Weight | TKW | Recorded after harvest. |

**Table S3.** Polymerase chain reaction (PCR) and agarose gel electrophoresis description.

<table>
<tr><th>Protocol/formulation</th><th>Description</th></tr>
<tr><td>Master mix composition</td><td>
<table>
<tr><td>Components</td><td>Volume (µl)</td></tr>
<tr><td>$H_2O$</td><td>15.9</td></tr>
<tr><td>PCR buffer (10x)</td><td>2.0</td></tr>
<tr><td>dNTP mix (10 Mm)</td><td>0.4</td></tr>
<tr><td>Forward primer (10 µM)</td><td>0.3</td></tr>
<tr><td>Reverse primer (10 µM)</td><td>0.3</td></tr>
<tr><td>Taq Polymerase (%U/µl)</td><td>0.1</td></tr>
<tr><td>DNA</td><td>1</td></tr>
<tr><td>Final volume</td><td>20</td></tr>
</table>
</td></tr>
<tr><td>PCR amplification regime</td><td>5 min at 94°C, 35x (30 s at 94°C, 30 s at 60°C, 60 s at 72°C) and 5 min at 72°C. The PCR was performed in a LifeTouch Thermal Cycler (Biozym Scientific GmbH, Hess. Oldendorf, Germany).</td></tr>
<tr><td>Agarose electrophoresis</td><td>From each sample, 6 µl were mixed with 2 µl of loading buffer, and from the resulting mix, 2 µl were loaded for electrophoresis. Water was used as negative control. Gels (3.0% agarose) were run for one hour at 100 V and 400 Amp. After electrophoresis, the agarose gels were visualized in a UV transilluminator using a Bio-Rad Laboratories gel-imaging system. The images were analysed in order to determine the PCR products size by comparison with the DNA size ladder and to classify the samples according to their genotypic classes.</td></tr>
</table>

**Table S4.** Allele and genotype nomenclature used in this study.

| Gene name | Location | Allele name | | Allele description | | Variant type | Aminoacid change |
|---|---|---|---|---|---|---|---|
| | | PI-614889 | CHEN-109 | PI-614889 | CHEN-109 | | |
| *HEN2* | chr4_56438962 | $A_1$ | $A_2$ | *A* | *T* | Intronic SNP | - |
| *SUF4* | chr12_80423460 | $B_1$ | $B_2$ | *A* | *G* | 3’ prime UTR SNP | - |
| *RUG3* | chr12_80551460 | $C_1$ | $C_2$ | *C* | *T* | 3’ prime UTR SNP | - |
| *RUG3* | chr12_80552842 | $D_1$ | $D_2$ | *C* | *T* | Missense SNP | p.Ala388Thr |
| *RUG3* | chr12_80553379 | $E_1$ | $E_2$ | *A* | *G* | Intronic SNP | - |
| *TKI1* | chr12_81633247 | $F_1$ | $F_2$ | *A* | *T* | Intronic SNP | - |
| *WRKY13* | chr12_81728382 | $G_1$ | $G_2$ | *C* | *A* | Missense SNP | p.Glu117Asp |
| *ATHB-15/CORONA* | chr4_51023634 | $H_1$ | $H_2$ | *G* | *A* | Missense SNP | p.Asp22Asn |
| *HEN2* | chr4_56441565 | $I_1$ | $I_2$ | *T* | *C* | Missense SNP | p.Ile724Met |
| *NPF2.6* | chr4_ 53557989 | $J_1$ | $J_2$ | *C* | *T* | Missense SNP | p.Ala93Val |
| *ERF113* | chr12_81516821 | $K_1$ | $K_2$ | *G* | *A* | Missense SNP | p.Pro326Leu |
| *FLD* | chr4_56844370 | $L_1$ | $L_2$ | *A* | *AT* | 3’ UTR InDel | - |
| *FLD* | chr4_56847562 | $M_1$ | $M_2$ | *TGAAC* | *T* | Intronic InDel | - |
| *TKI1* | chr12_81633685 | $N_1$ | $N_2$ | *T* | *A* | Missense SNP | p.Gln445Leu |
| *MET1b* | chr4_56534732 | $O_1$ | $O_2$ | *ATT* | *A* | Frameshift | p.Gln222fs |
| *MET1b* | chr4_56534915 | $P_1$ | $P_2$ | *AGTT* | *A* | Disruptive Inframe Deletion | p.Lys161_Leu162delinsMet |
| *RICESLEEPER3* | chr4_55091902 | $Q_1$ | $Q_2$ | *A* | *AATTCCT* | Disruptive Inframe Insertion | p.Ile281_Thr282insProIle |

**Table S5.** Genetic and phenotypic segregation for two traits in the $F_2$ and $F_3$ populations. Red axil pigmentation was determined five weeks after sowing. The InDel marker JAASS5 was described by Zhang et al. (2017). $R_1$ and $R_2$ represent the 189 bp and 164 bp alleles, respectively.

| Plants | **$F_2$ population** | | | | | | | **$F_3$ population** | | | |
|---|---|---|---|---|---|---|---|---|---|---|---|
| | Red axil pigmentation | | $\chi^{2\,a}$ | JASS5 genotype | | | $\chi^{2\,b}$ | JASS5 genotype | | | $\chi^{2\,c}$ |
| | Red | Green | | $R_1R_1$ | $R_1R_2$ | $R_2R_2$ | | $R_1R_1$ | $R_1R_2$ | $R_2R_2$ | |
| O | 254 | 82 | 0.06 | 8 | 28 | 12 | 2.00 | 58 | 57 | 79 | 5.02 |
| E | 252 | 84 | | 12 | 24 | 12 | | 72.75 | 48.50 | 72.75 | |

E: expected, O: observed.
[a] 3:1 segregation, $\chi^2_{(0.95;1)} = 3.84$
[b] 1:2:1 segregation, $\chi^2_{(0.95;2)} = 5.99$
[c] 3:2:3 segregation, $\chi^2_{(0.95;2)} = 5.99$
